# Biosynthesis of minimal C-phycocyanin chromophore assemblies in *E. coli* provides a platform to dissect protein-mediated tuning of exciton transfer

**DOI:** 10.1101/2025.10.13.682215

**Authors:** Deborah L. Zhuang, Derrick S. Chuang, Annika B. Velez, Kylee B. Gao, Sophia M. Velilla, Mia Sweeny, Jeffrey A. Chen, Krishna K. Niyogi, Masakazu Iwai, Matthew B. Francis

**Author notes:** **Corresponding Author** Matthew B. Francis.

## Abstract

Cyanobacteria are arguably among the most evolutionarily successful organisms on Earth, inhabiting a wide range of ocean, fresh water, soil, and even desert environments on every continent. The cyanobacterial phycobilisome consists of stacks of disk-like light-collecting moieties, allophycocyanin (APC) and phycocyanin (CPC), with covalently bound phycocyanobilin (PCB) pigments. The ways in which the energies of the specific chromophores in these complexes are tuned by the protein to achieve its highly efficient and directional energy transfer are not fully understood, as complex combinations of decay pathways are occurring simultaneously and competitively through this elaborate light-harvesting system. This makes it difficult to extract information about isolated protein-pigment interactions. We provide herein a description of a useful new experimental platform in which we have recombinantly expressed a fully functioning CPC complex and selectively created minimal chromophore sets to study their individual contributions to the overall CPC spectra. Structural and computational analysis of this protein system have provided a greater understanding of how the protein environment serves to alter the photophysics of each of these chromophores. Introduction of a quencher into various positions within CPC confirmed the ability of the protein environment to tune the directionality of energy transport in this assembly. Further mutational analysis suggested the roles of key amino acids surrounding the chromophores, showcasing the utility of heterologous expression techniques for understanding the effects of structure on EET mechanisms in the phycobilisome.

## INTRODUCTION

Given the rich and central biological history of cyanobacteria, researchers have long been motivated to study the photosynthetic systems within these organisms. In them, light is collected as photons and transferred as excitons to the chlorophyll-based photosystems by an elaborate light-harvesting structure known as the phycobilisome. [1, 2, 3] The cyanobacterial phycobilisome comprises two principal light-collecting moieties within its structure; it has a central core made up of allophycocyanin (APC) disk-like protein-pigment complexes that are surrounded by rods containing phycocyanin (CPC) disks. These components are held together by a series of linker proteins that do not contain any bound pigments, but are hypothesized to aid in the directionality of energy transfer in these megastructures. [4, 5] For both CPC and APC, the assemblies consist of alpha and beta subunits arranged into single-layer α_3_β_3_ disks that serve as the basic phycobilisome building blocks (**Figure 1a**). Both the α- and β-subunits are covalently modified with the same linear tetrapyrrole chromophore, phycocyanobilin (PCB), which is covalently attached to specific cysteine residues within the protein subunits through thioether linkages.

**Figure 1.**
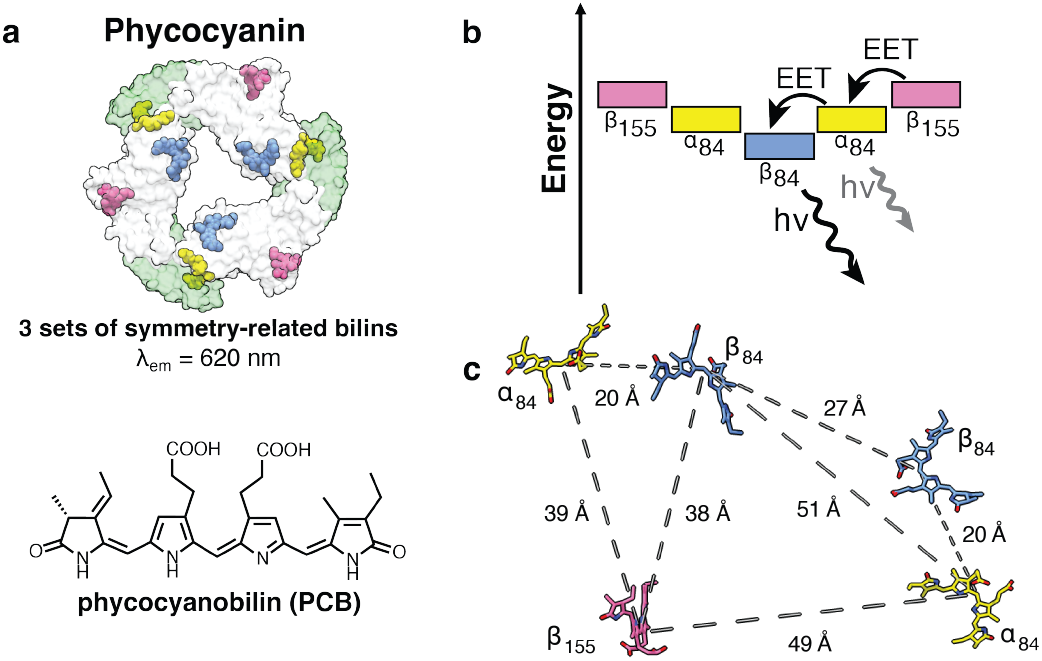
Structure of the phycocyanin (CPC) assembly from *Synechococcus elongatus* PCC 7942. **(a)** The αβ assembly contains three sets of chemically identical phycocyanobilin (PCB) chromophores in unique protein environments, leading to their distinctive spectral properties. **(b)** As reported, the environmental differences of each set of chromophores lead to a well-described energy flow from the peripheral β_155_ PCB towards the terminal emitter, the β_84_ PCB. **(c)** When mapping the possible energy transfer pathways in CPC, there are many combinations of PCB interactions that are within 50 Å of each other, leading to a highly interconnected chromophore network that makes extracting distinct contributions of the individual environments difficult.

The prevailing hypothesis for phycobilisome function, including that for the CPC complex, involves the absorption of light energy in the periphery of the assembly to the core of the structure. The excitons are ultimately carried to the chlorophylls of membrane-bound photosystems (**Figure 1b**). [6] The ways in which the energies of the specific chromophores are tuned by the proteins of the system to achieve this are not fully understood. Early time-resolved spectroscopic examinations of fully intact CPC structures isolated from native organisms have revealed estimates for excitation energy transfer (EET) rates between individual chromophores, and these have been matched to the distance relationships within the assemblies using Förster energy transfer theory. [7, 8] While this has provided insight into the systems as they occur in nature, complex combinations of decay pathways are occurring simultaneously and competitively and thus are difficult to differentiate in fully intact systems. For example, a complex network of 5 possible interactions is possible if all chromophore distances below 50 Å are considered (**Figure 1c**). Moreover, it is difficult to match the subtly different (but likely very important) excitation and absorption spectra of the individual pigments to their protein environments using this aggregate approach.

To obtain isolated EET characteristics for the different chromophores, we prepared a set of CPC variants that systematically lack each set of PCB chromophores using a heterologous, modular expression system in *E. coli* (**Figure 2a**). This was achieved by controlling the expression of the lyase enzymes responsible for regiospecifically modifying each position within CPC. Thus, by systematically excluding or including individual chromophore attachment sites (α_84_, β_84_, β_155_), we achieved controlled formation of partially or fully chromophorylated CPC variants. Such an approach allows the spectroscopic properties (e.g. absorption/emission spectral shifts and fluorescence lifetime modulations) specific to each chromophore subset to be determined unambiguously. Moreover, it substantially limits the number of available EET pathways, providing accurate decay values for single chromophores and defined pairs. In this work, we apply biochemical, spectroscopic, and computational tools to correlate the structural features of the microenvironments that surround the chromophores to key electronic transitions, thus obtaining a clearer understanding of the individual behaviors of the different pigment molecules that allow vectoral EET to proceed. All these data are matched to those of fully intact CPC assemblies, both from recombinant expression in *E. coli* as well as those isolated from the native cyanobacterium. This approach yields a more accurate understanding of the spectroscopic differences between individual chromophores. Some of the CPC mutants identified using heterologous expression were expressed in their native hosts, allowing the effects on viability and growth to be determined. Specifically, we found that the β_84_ chromophore plays a key role in proper phycobilisome function; in contrast, the α_84_ and β_155_ chromophores are not required for growth under low and standard light conditions but are necessary under high light conditions. This suggests they could play a role in photoprotection under high photon flux.

**Figure 2.**
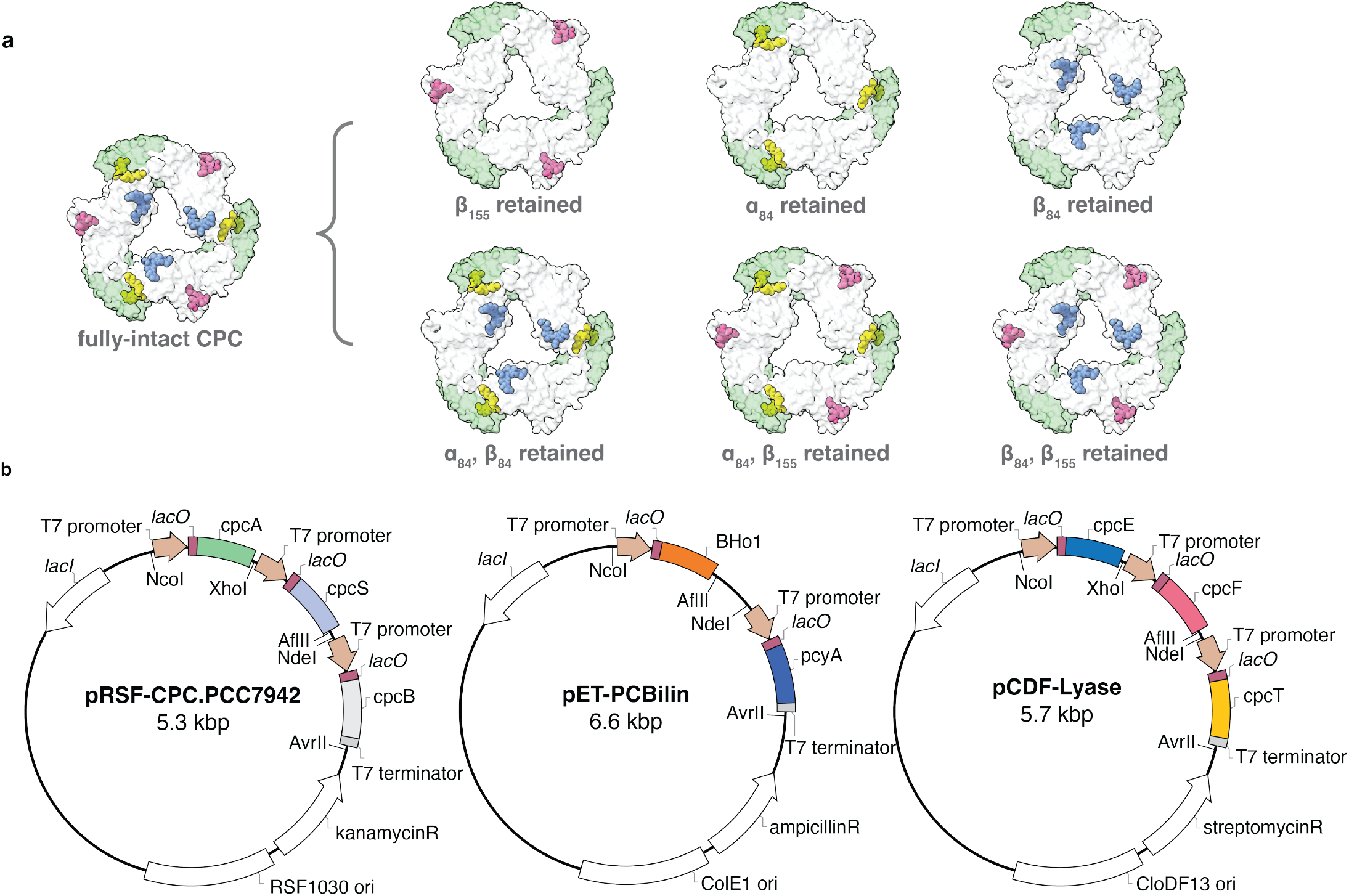
Expression system for recombinant production of CPC in *E. coli*. **(a)** A series of CPC disks containing single PCB sets were prepared to obtain accurate and unambiguous spectral contributions of each protein environment. To add pathways and introduce coupling interactions, disks with two sets of PCB pigments were also generated. **(b)** These chromophore knockouts were performed by removing the genes encoding the lyases responsible for regiospecifically attaching PCBs to specific cysteine residues. A three-plasmid expression system with orthogonal antibiotic resistance markers and origins of replication was used to do this.

The results provided by the native organism highlight the benefit of the recombinant expression system in *E. coli*, where the ability of the organism to propagate is no longer dependent on fully “normal” function of the proteins. This allows for these minimal sets to be produced readily for multimodal characterization with greater speed, flexibility, and reliability. The directionality of energy transfer across the recombinant CPC assembly was then investigated by attaching a small-molecule quencher to unmodified cysteine residues within each reduced-chromophore CPC to compare the fluorescence lifetimes with and without quenching at the neighboring positions. These results show the contribution of each pathway to the full network of EET processes occurring within the fully intact CPC and suggest that the protein plays a role in funneling the excitons towards the terminal emitter at β_84_. The flexibility of this heterologous expression system allows us to learn how specific protein environments shape the spectroscopic properties of individual chromophores, and it shows which energy transfer pathways dominate as excitons move through protein assemblies as different sets of chromophores are introduced. Taken together, these experiments provide great detail and accuracy of how proteins tune the spectral properties of embedded chromophores.

## RESULTS AND DISCUSSION

Existing reports have described heterologous expression platforms to produce various components of the PBS to study *in vitro*. Multi-plasmid expression systems have been constructed to produce highly chromophorylated APCs, which require only a single CpcS lyase enzyme to modify both sites of chromophore attachment. [9] This has shed light on the structural basis of certain far-red light absorbing variants of APC, illuminating features in those APC paralogs that produce drastic redshifts in absorbance. [10] However, this recombinant platform has yet to be fully optimized to study CPC due to the three sites of chromophore attachment requiring multiple lyase enzymes. [11, 12, 13] There exist studies constructing α- or β-subunits of chromophorylated CPC or partially chromophorylated CPC to study their antioxidant or radical scavenging activities. [14, 15, 16] Efforts have also been made to improve chromophorylation efficiency through metabolic engineering of the *E. coli* hosts and to evolve the lyases to improve chromophorylation. [17] In general, these systems were not adaptable for the studies described herein, therefore requiring a new expression platform that allowed greater control over the chromophore content of the proteins.

### Construction of a phycocyanin expression system in *E. coli*

We derived our sequences for the CPC α and β chains from the cyanobacterial species *Synechococcus elongatus* sp. PCC 7942 due to the high-resolution crystal structures of the phycobilisome (PBS) components in this organism. [18] To construct a fully intact CPC expression system for these proteins in *E. coli*, we used a set of three orthogonal Duet^TM^ vectors that contain two T7 promoter regions (**Figure 2b**), each being under a different origin of replication. We inserted genes encoding the α and β CPC subunits along with the gene encoding the CpcS lyase (preceded by an additional T7 promoter and ribosome binding site) into a high copy number pRSFDuet^TM^-1. Into the pETDuet^TM^-1 vector, we inserted the genes that convert free heme within *E. coli* to PCB. This is achieved by converting hemin into biliverdin-IXα by heme oxygenase 1 from the thermophilic cyanobacterial species *Thermosynechococcus elongatus* BP-1 (BHo1) for greater thermal stability, which then undergoes a 4-electron reduction with phycocyanobilin:ferredoxin oxidoreductase (PcyA). This biosynthetic pathway to constructing PCB in *E. coli* hosts is depicted in the Supplementary Information. Finally, into pCDFDuet^TM^-1 we inserted the genes encoding for additional lyases: the heterodimer CpcE/CpcF and monomeric CpcT. The sequence specificity for each lyase enzyme to modify cysteine positions with PCB has been reported, where the β_155_ cysteine is modified by CpcT, β_84_ is modified by CpcS, and α_84_ is modified by the dimer of CpcE/CpcF. [11, 12, 13] Therefore, by excising the genes that encode for different lyase enzymes, we can control which cysteine positions are and are not modified with the chromophore. This strategy leaves the native cysteine in unchromophorylated pockets to preserve their amino acid composition.

Following expression and lysis, this material was pelleted via ammonium sulfate precipitation, collected, and dialyzed against a low-salt buffer (20 mM triethanolamine, pH 7.2) before purification with a DEAE anion exchange column with a 0-600 mM NaCl gradient. The first peak containing CPC was collected before buffer exchanging into a high ionic strength buffer of 200 mM sodium acetate at pH 5.5, resulting in pure CPC trimer assemblies. This was confirmed with ESI-MS to determine both purity and the degree of chromophore modification (**Figure 3a**). Native gel electrophoresis (**Figure 3b**) showed only one intense fluorescent band at roughly 120 kDa corresponding to the trimeric assembly. From these characterization data, we concluded the the minimal CPC constructs expressed using this platform form homogenous assemblies with covalently attached PCB chromophores, which are important features for recapitulating the properties of native CPCs.

**Figure 3.**
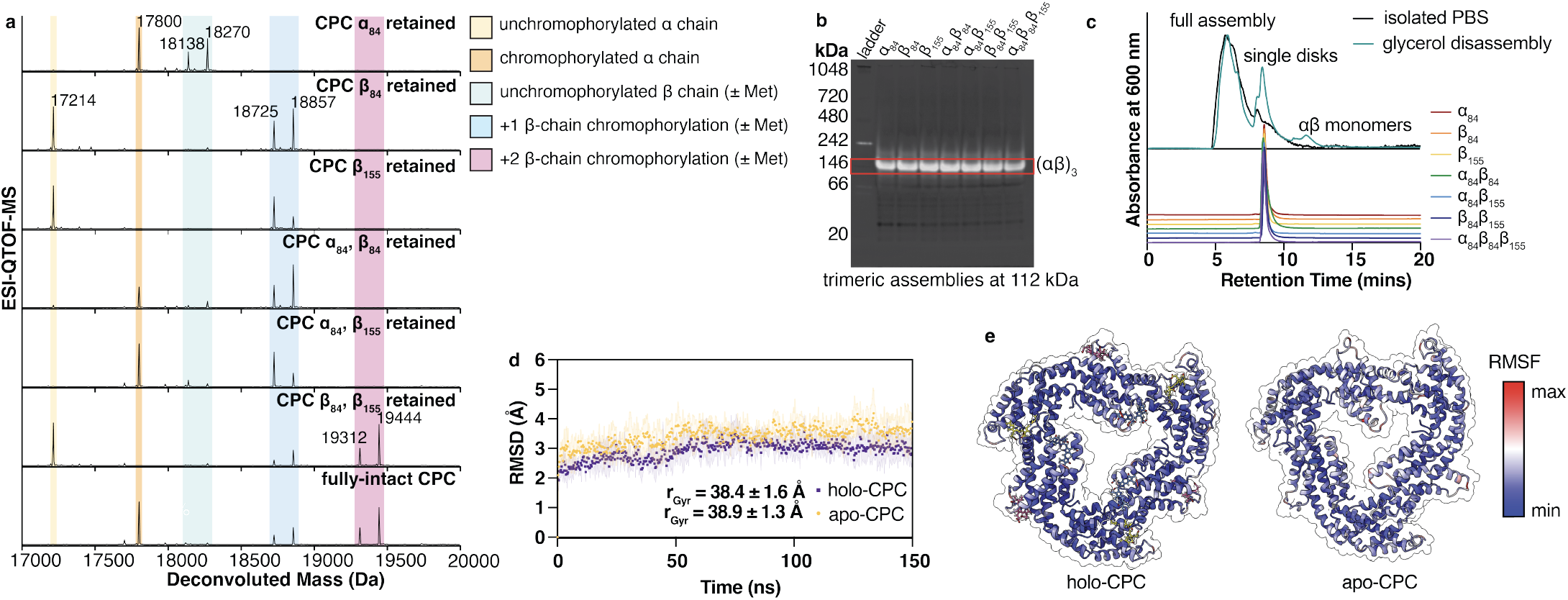
Characterization of recombinantly expressed phycocyanin variants. **(a)** ESI-MS showed successful chromophorylation for each variant. **(b)** Native PAGE showed a single brightly fluorescent band overlaid with Coomassie blue staining, indicating a trimeric assembly state around 112 kDa. **(c)** Normalized peaks from size-exclusion HPLC showed clean assembly states matching the elution time of the trimeric, single disk phycocyanin complexes taken from disassembling the full phycobilisome isolated from *S. elongatus* PCC 7942. **(d)** Triplicate MD simulations show no significant differences in the average protein backbone RMSD or the average radius of gyration when comparing the holo- and apo-CPC assemblies. This suggests that the protein does not collapse in the absence of chromophores. **(e)** From the trajectories, we compared frames of the holo-(left) and apo-(right) CPC colored by the per-residue RMSF to find that even with greater fluctuation in the protein α-carbons near the sites of chromophore deletion, the overall assembly state remains intact.

The elution times from size-exclusion HPLC for the recombinantly expressed CPCs were compared to the structures isolated from the wild type *S. elongatus* PCC 7942 (**Figure 3c**). The recombinant CPCs matched exactly with the retention time of single disk α_3_β_3_ CPC isolated from the native PCC 7942 organism following 50% (v/v) glycerol-promoted disassembly from the phycobilisome. [19] In a negative control where no lyases were expressed, this resulted in no chromophorylation of the α- or β-chains, and as a result, we did not observe high-intensity fluorescence (Figure S1). This shows that there is minimal nonspecific chromophore binding in the absence of the proper lyases. As has been observed previously, that there is minimal nonspecific chromophore binding in the absence of the proper lyases. As has been observed previously, the free PCB chromophore alone is not appreciably fluorescent. [11] Furthermore, the lyases themselves do not contribute to the overall fluorescence signal when mutations to the chromophore binding sites prevented the covalent addition of the chromophore (Figure S2).

Molecular dynamics (MD) simulations were performed in triplicate on trimeric CPC protein assemblies to compare fully-chromophorylated systems with the chromophore knockouts. These simulations showed that the overall protein backbone and the chromophore-binding cavities remained stable in the absence of covalently bound PCB over the course of 150 ns. In addition, negligible collapse of the chromophore-binding cavities was observed (**Figure 3d,e**). From these simulations, we were then also able to calculate the solvent-exposed surface area (SASA) of each chromophore (Table 1), in which we observed a linear relationship between the SASA and the Stokes shift. The cosine content of the first principal component of the trajectory indicated that adequate sampling had been used (further described in the Supplementary Information). [20, 21]

**Table 1.**
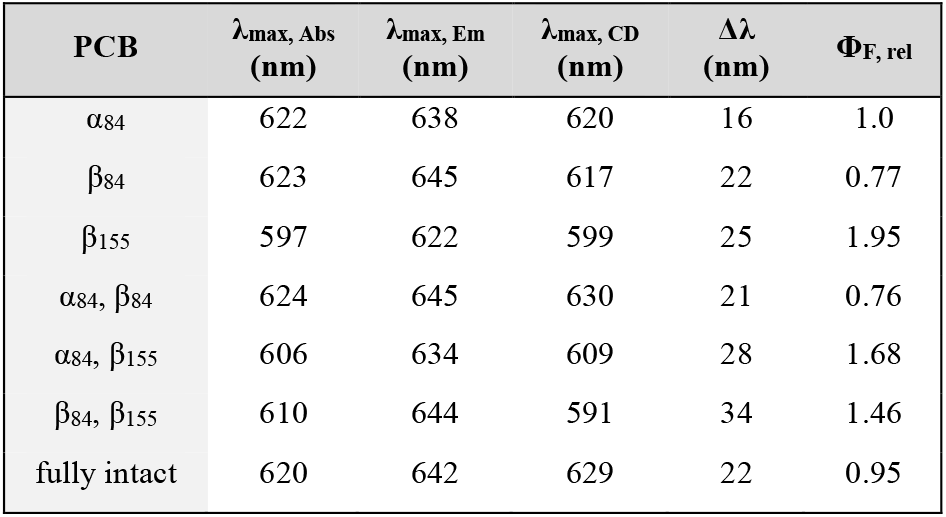
Experimentally measured spectral data for each of the CPC chromophore sets.

| PCB | $\lambda_{\text{max, Abs}}$<br>(nm) | $\lambda_{\text{max, Em}}$<br>(nm) | $\lambda_{\text{max, CD}}$<br>(nm) | $\Delta\lambda$<br>(nm) | $\Phi_{\text{F, rel}}$ |
| --- | --- | --- | --- | --- | --- |
| $\alpha_{84}$ | 622 | 638 | 620 | 16 | 1.0 |
| $\beta_{84}$ | 623 | 645 | 617 | 22 | 0.77 |
| $\beta_{155}$ | 597 | 622 | 599 | 25 | 1.95 |
| $\alpha_{84}, \beta_{84}$ | 624 | 645 | 630 | 21 | 0.76 |
| $\alpha_{84}, \beta_{155}$ | 606 | 634 | 609 | 28 | 1.68 |
| $\beta_{84}, \beta_{155}$ | 610 | 644 | 591 | 34 | 1.46 |
| fully intact | 620 | 642 | 629 | 22 | 0.95 |

### Spectroscopic differences between chromophores in each unique phycocyanin environment

Having characterized the recombinantly expressed phycocyanin that assembles into a single population of CPC trimers, we then pursued the spectral characterization of each variant. In general, these spectra matched well with the reported values for assembled CPC complexes. [22, 23, 24] Despite chemically identical chromophores being ligated to each group of cysteine sites within the CPC protein, the steady-state absorbance and fluorescence emission measurements showed distinct spectral profiles for each chromophore, resulting from its unique configuration and protein environment. These data are summarized in **Figure 4** and tabulated in Table 1. As reported, the outermost β_155_ chromophore is at the highest energy, showing the shortest wavelength absorbance and emission maxima, followed by α_84_, then β_84_ at the lowest energy.

**Figure 4.**
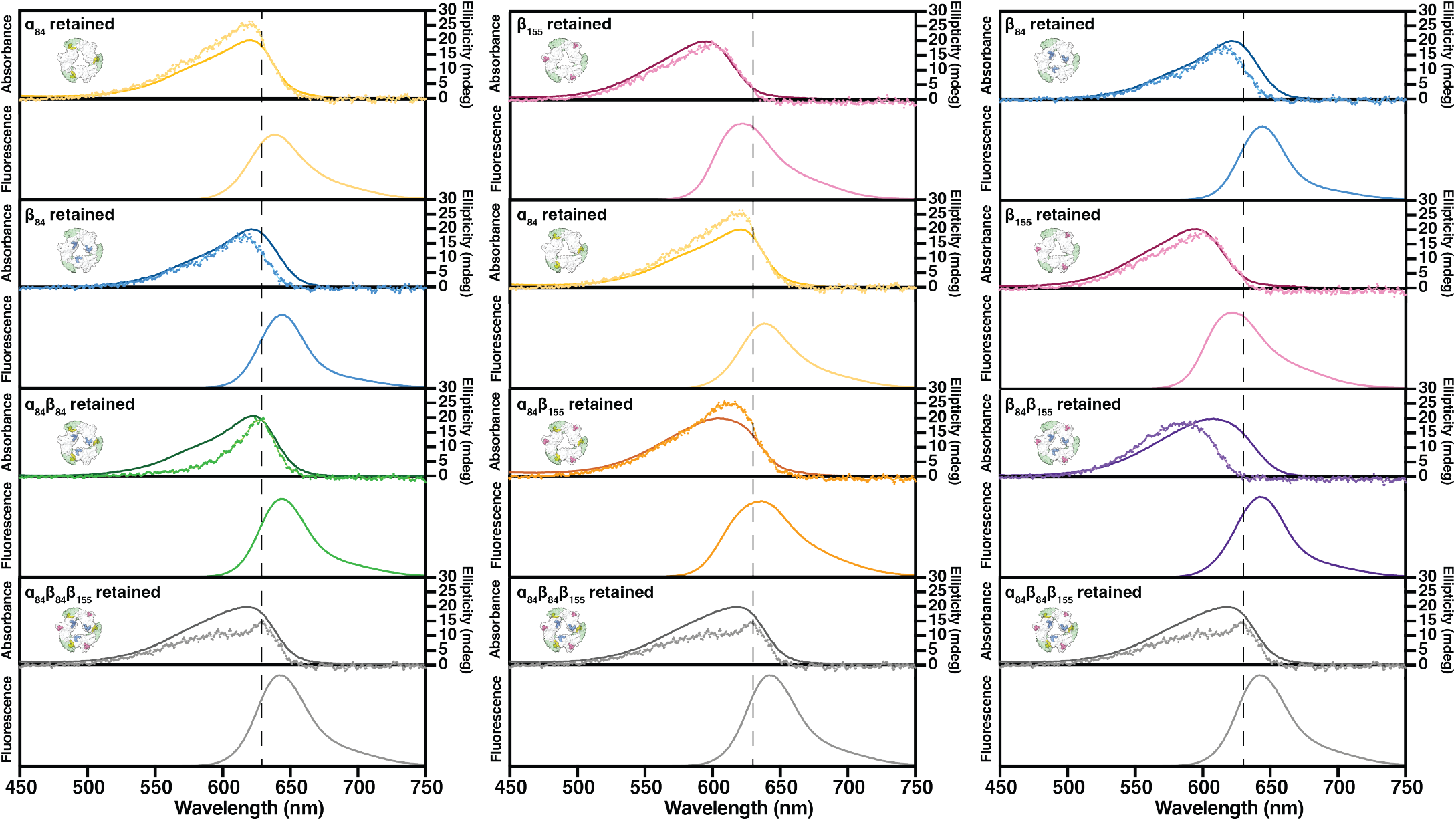
Normalized steady-state spectra of each CPC chromophore combination. Absorbance profiles showed differing energy levels for each chromophore, with the chromophore at β_155_ having the highest energy. The double chromophore variants showed broader spectra resulting from the contributions of each pigment. The CD spectra of the chromophores (dotted lines) were similarly shaped to the absorption bands (solid lines). However, red-shifted maxima were observed in the CD spectra due to excitonic coupling between α_84_/β_84_ chromophore pairs (green and grey traces). The emission spectra of the fully intact CPC (grey) showed little contribution from the β_155_ emission (pink).

For the purposes of comparison, TD-DFT was used to produce spectral predictions for each of these chromophores as taken from the crystal structure. The calculated spectra were well matched to the experimentally obtained spectra (Figure S5). The CD spectra of each of these chromophores in the visible light range showed similar bandshapes to the absorption spectra, but sharp, red-shifted maxima appeared whenever the α_84_/β_84_ chromophores in the monomer-monomer interface were both present and able to interact (**Figure 4**). These features have been assigned to the excitonic coupling interactions between the α_84_/β_84_ chromophores. [25, 26, 27] Taken together, these results suggest that the heterologously expressed chromophore systems provide accurate information about the spectroscopic properties of the individual chromophores and their aggregate combinations.

### Impact of chromophore deletions on cyanobacterial growth rates under varying light conditions

In addition to the recombinant CPC mutants we examined *in vitro*, we also tested the impact of two PCB deletions in CPC *in vivo* using *Synechococcus elongatus* PCC 7942. We generated the two mutant lines through site-directed mutagenesis of the three chromophore-binding cysteine residues (α-85, β-83, β-154, equivalent to positions α-84, β-84, and β-155 in the recombinantly expressed sequences) to modify pigment binding in the CPC. Because we could not knock out the lyases in the wild-type strains without deleterious effects to other components in PBS, we instead changed the cysteine residues to valines that could not be modified with PCBs. The first mutant line (Δα_85_Δβ_154_) has C-to-V mutations at the α-85 and β-154, leaving only the interior β-83 PCB site intact, while the other mutant line (Δα_85_Δβ_83_) has C-to-V mutations at α-85 and β-83, retaining only the outermost β_154_ PCB site.

The photosynthetic growth rates of these two mutant lines were then compared with those of the wild type (WT). Under low light conditions (LL; ∼40 μmol photons m^-2^ s^-1^), the Δα_85_Δβ_154_ line showed a similar growth rate to the WT, but the Δα_85_Δβ_83_ line had a drastically hindered growth rate (**Figure 5a**). This observation was consistent with previous studies, suggesting that the β_83_ PCB serves as the primary acceptor that transfers energy down the central channel of CPC rods to the APC phycobilisome core. [28, 29] As expected, the growth under dim light conditions (DL; ∼3 µmol photons m^-2^ s^-1^) caused a much slower rate even for WT, but the Δα_85_Δβ_154_ line still showed a similar rate to the WT (**Figure 5b**), signifying the importance of the interior β_83_ PCB for proper energy transfer and the photosynthetic growth. Interestingly, however, the Δα_85_Δβ_154_ line showed a severe reduction in growth under high light conditions (HL; ∼285 µmol photons m^-2^ s^-1^), even though the WT grows normally (**Figure 5c**). This might indicate that these missing PCBs at α_85_ and β_154_ provide energy transfer pathways that are necessary under HL, directing an excess amount of excitation energy to minimize photodamage. It has been known that excess energy can be dissipated in the CPC in the phycobilisome of *S. elongatus* PCC 7942 through an exciton-exciton annihilation mechanism within its rods before the energy is transferred downhill to the APC. [30] Because the orange carotenoid protein (OCP), which causes the nonphotochemical quenching of excitation energy, is absent in *S. elongatus* PCC 7942, the quenching mechanisms in the PBS are not well-understood. [31]

**Figure 5.**
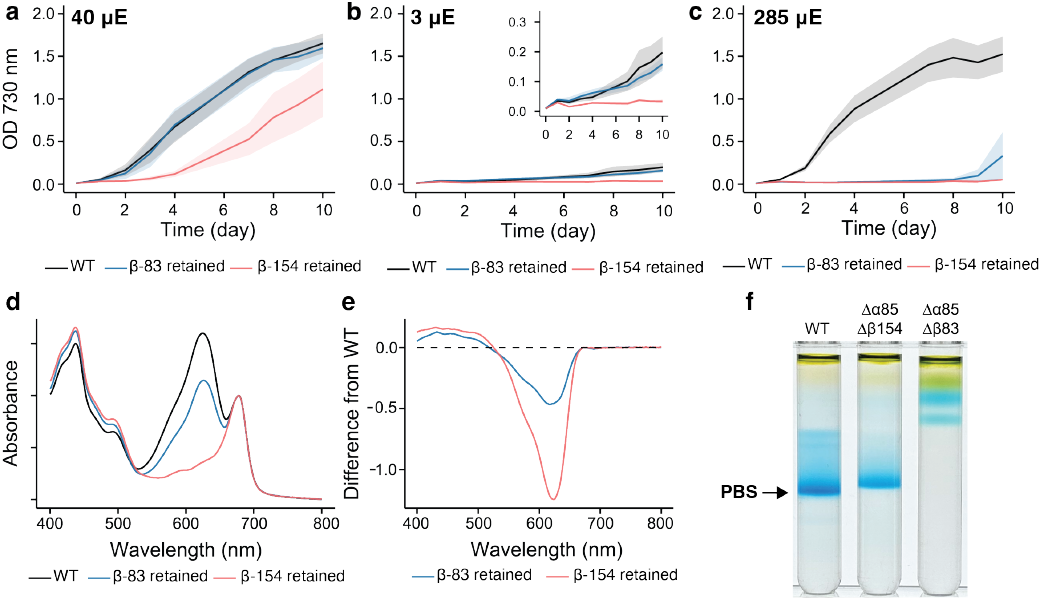
The impact of PCB deletion on photosynthetic growth and structural stability of the PBS. Photosynthetic growth comparison between the WT and two mutants (Δα_85_Δβ_154_, retaining β_83_ and Δα_85_Δβ_83_, retaining β_154_) of *S. elongatus* PCC 7942 under three different light intensities: **(a)** ∼40 (LL), **(b)** ∼3 (DL), and **(c)** ∼285 (HL) µmol photons m^-2^ s^-1^ (µE). **(d)** The absorbance spectra are shown for the cyanobacterial cultures of WT and the two mutant strains. The data were normalized at the chlorophyll *a* peak in the Q_y_ region. **(e)** Difference spectra between WT and the two mutants are shown. **(f)** Sucrose gradients separating the intact and disassembled PBS are shown for the WT and the mutants.

To examine how the C-to-V mutations at the α_85_, β_83_, and β_154_ sites affect the light absorption, we measured absorbance spectra using the grown culture of each line. Intriguingly, the Δα_85_Δβ_154_ line, which has a similar growth rate to the WT under LL and DL, shows approximately 35% reduction of the PBS absorption as compared to the WT (**Figure 5d,e**). This result is aligned with the growth phenotype data, in that the PCBs at the α_85_ and β_154_ sites have accessory roles in light absorption under LL but become critical to manage the excess amount of excitation energy under HL conditions. On the other hand, the Δα_85_Δβ_83_ line, which already shows a hindered growth rate under LL, completely lacks the PBS absorption (**Figure 5d**). To determine the factors affecting PBS absorption, we performed purification of the PBS using sucrose gradient ultracentrifugation. Surprisingly, both C-to-V mutations impact the structural stability of PBS. The Δα_85_Δβ_154_ line contains a slightly smaller PBS than the WT (**Figure 5f**), indicating that the PCBs at the α_85_ and β_154_ sites are not just accessory chromophores but also necessary for the stable formation of the PBS structure *in vivo*. On the contrary, the Δα_85_Δβ_83_ line lacks the PBS structure, which is aligned with the observation of the absence of its absorption. Although we cannot rule out the possibility that these different formations of the PBS structures occurred during the sample preparation or due to changes in protein expression of adjacent linker proteins caused by the mutation in CPCs, our observation suggests that the PCBs play roles not only in excitation energy transfer within the PBS structure but also in its stability *in vivo*.

These results underscore the utility of preparing PBS sub-components through heterologous expression as a useful complement to *in vivo* methods. Because they can be obtained independently from cyanobacterial survival, they allow the optical effects of chromophore changes to be distinguished from larger phenotypic differences observed using the *in vivo* systems. They are also amenable to studying large numbers of mutants with greater throughput.

### Tracking energy flow in the phycocyanin assembly

It is well-appreciated that the protein environment itself in CPC serves some role in directing the flow of excitation energy across the complex. As previously discussed, the energy levels of the chromophores are thought to direct absorbed light energy from β_155_ to the terminal emitter at β_84_. This pathway has been elegantly explored through multiparametric single-molecule photophysical measurements of CPC and APC, relying on step-wise photobleaching of each chromophore to characterize individual pigment sites. [32, 33] As a useful way to complement these studies and confirm this pathway, our method provides a unique opportunity to measure the lifetimes of the individual chromophores directly with unambiguous assignments of the contributing chromophores, as well as the aggregate lifetimes of all pairwise combinations. These data are summarized in Table 2. From these measurements, we see that the longer lifetime of β_155_ is unperturbed upon adding the chromophore at position α_84_, but it shortens when β_84_ is present. This parallels the quick transfer of energy from β_155_ to β_84_ (*k* ≈ 50 ps) and slow transfer from β_155_ to α_84_ as predicted from a FRET model based on the relative geometry of the chromophores (calculated in Table S2). [7, 22, 26]

**Table 2.**
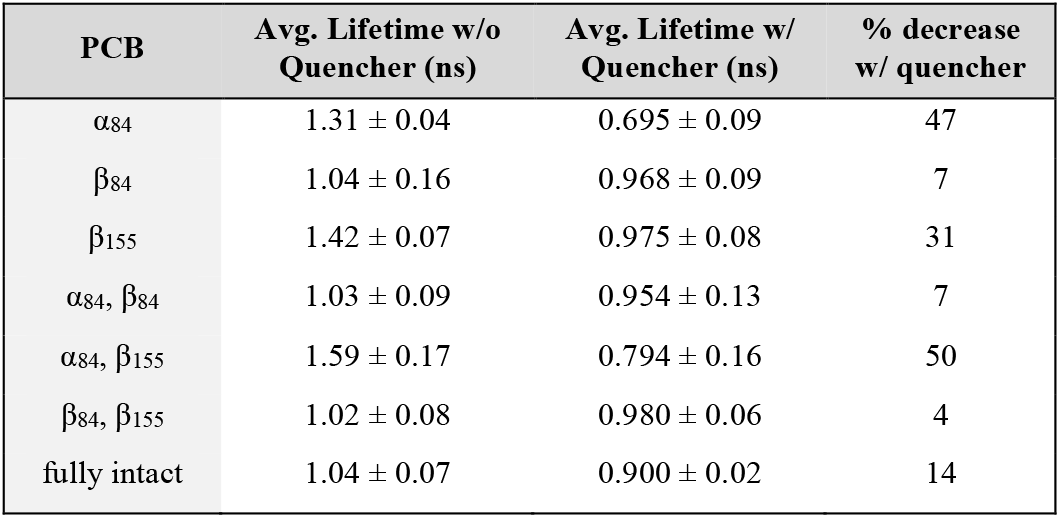
Amplitude-averaged fluorescence lifetime measurements for each set of CPC chromophore variants.

| PCB | Avg. Lifetime w/o Quencher (ns) | Avg. Lifetime w/ Quencher (ns) | % decrease w/ quencher |
| --- | --- | --- | --- |
| $\alpha_{84}$ | $1.31 \pm 0.04$ | $0.695 \pm 0.09$ | 47 |
| $\beta_{84}$ | $1.04 \pm 0.16$ | $0.968 \pm 0.09$ | 7 |
| $\beta_{155}$ | $1.42 \pm 0.07$ | $0.975 \pm 0.08$ | 31 |
| $\alpha_{84}, \beta_{84}$ | $1.03 \pm 0.09$ | $0.954 \pm 0.13$ | 7 |
| $\alpha_{84}, \beta_{155}$ | $1.59 \pm 0.17$ | $0.794 \pm 0.16$ | 50 |
| $\beta_{84}, \beta_{155}$ | $1.02 \pm 0.08$ | $0.980 \pm 0.06$ | 4 |
| fully intact | $1.04 \pm 0.07$ | $0.900 \pm 0.02$ | 14 |

As an additional experimental examination of the contribution of each potential transfer pathway to the fully intact CPC assembly, we took advantage of the unmodified cysteine residues remaining in the chromophore subsets. To each of the remaining cysteine positions within each CPC variant, we conjugated a maleimide-functionalized methylene blue dye (**Figure 6a-c**, Atto MB2, λ_max, abs_ = 668 nm, Φ_F_ < 0.04) that exhibits appreciable spectral overlap with each PCB chromophore (Figure S6, S7). We then measured the changes in the fluorescence lifetimes of the complexes as dynamic quenching occurred (**Figure 6d**). As excitation energy absorbed by PCB pigments is transferred to the quenchers, a decrease in the complex’s fluorescence lifetime is expected if the energy transfer rate from PCB to the quencher is comparable to or faster than the fluorescence lifetime of the PCB. This was indeed observed, and the data are listed in Table 2. We can envision that the transfer efficiency from chromophore to quencher in this system will not be fully efficient due to alignment changes of the respective transition dipole moments (as shown in ligand docking experiments *in silico* shown in Figure 6b and calculated in the Supplementary Information, Figure S8). However, through this method we can still compare the percent contribution of each transfer pathway by comparing the ratios of lifetime decrease.

**Figure 6.**
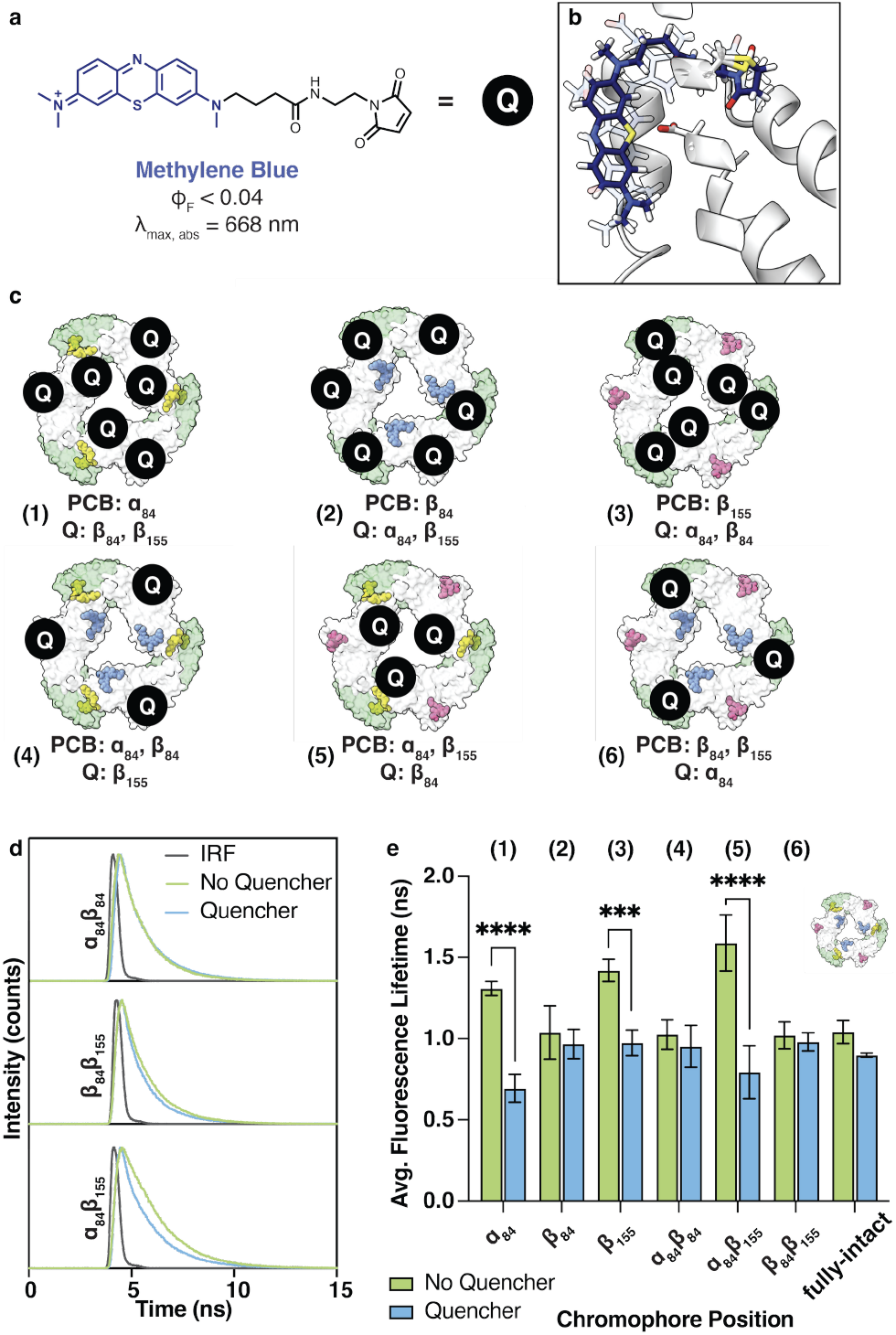
Studies of the directionality of energy transfer within CPC capitalizing on the free cysteines left in this expression system. **(a)** A methylene blue-based quencher was covalently attached via maleimide chemistry to the unmodified cysteines in the CPC minimal sets to examine the positional effects on the extent of exciton quenching. **(b)** From docking experiments, these quenchers occupy the same binding site as the PCB chromophores, resulting in similar alignments of their transition dipole moments. **(c)** The extent of quenching can be measured at various positions within the CPC assembly to determine the directional flow of energy transfer across the complex. **(d)** Time-resolved emission measurements were used to measure dynamic quenching processes. The faster rate of exciton transfer to quencher sites compared to fluorescence rates led to decreased fluorescence lifetimes of some CPC complexes. **(e)** The amplitude-weighted average lifetimes for each sample’s exponential decay were compared with and without the presence of the conjugated quencher. The errors given are those obtained from the variations between several independent preparations. The statistical errors of the decay analysis are smaller than these values. Samples that showed statistical significance (from an unpaired 1-tailed *t*-test) in fluorescence lifetime decreases are highlighted with asterisks; differences in other measurements were not statistically significant.

From Figure 6e, we see the largest decrease in lifetime for CPC complexes lacking the β_84_ chromophore (**1, 3**, and **5** in **Figure 6e**), where CPC complexes with chromophorylated α_84_, β_155_ show ∼50% decrease in fluorescence lifetime when the quencher is attached at β-84. This signifies that a majority of the excitons are transferred to the β-84 quencher before they can emit. Likewise, when only β_155_ is chromophorylated and quenchers are at position α-84 and β-84, there is a large 31% decrease in fluorescence lifetime. The lack of dynamic quenching when the quencher is positioned solely at β-155, as in the cases where α_84_, β_84_ are chromophorylated, or β_84_ alone is chromophorylated, suggests that the protein environment serves some role in the directional energy transfer in the CPC assembly, preventing the reversal in energy flow outwards towards the β-155 site. Despite their spectral similarity and fast transfer rates between the α_84_ and β_84_ positions, we interestingly observed that, when the quencher was positioned on α-84, there was no significant quenching measured even if there is a chromophore in the nearby β_84_ position. This seems to suggest that the energy transfer is largely tuned by the protein, such that there is no reversal in energy flow to the β_155_ or α_84_ position, and excitons are largely and rapidly funneled to position β_84_. The fully intact CPC with all three sets of coexpressed lyases was also used as a control in these quenching experiments. The incomplete chromophorylation of this system (**Figure 3a**), leaving free cysteine residues exposed, could account for a slight decrease in the measured lifetimes in these systems, though it was not statistically significant in these studies. Thus, the contribution of nonspecific quencher binding to the lifetime measurements is minimal.

When we look at key interactions in the crystal structure that could serve to tune the flow of excitons in CPC, a standout feature is the network of aromatic residues surrounding the coupled α_84_/β_84_ chromophore pair (**Figure 7a**). These chromophores show a similarly positioned tyrosine residue interacting with a highly conserved aspartate; it is posited that this tyrosine residue stabilizes the chromophore in the binding pocket by restricting its range of motion. Likewise for each chromophore, there is a conserved aromatic network with π-stacking properties that could be further explored through targeted mutagenesis.

**Figure 7.**
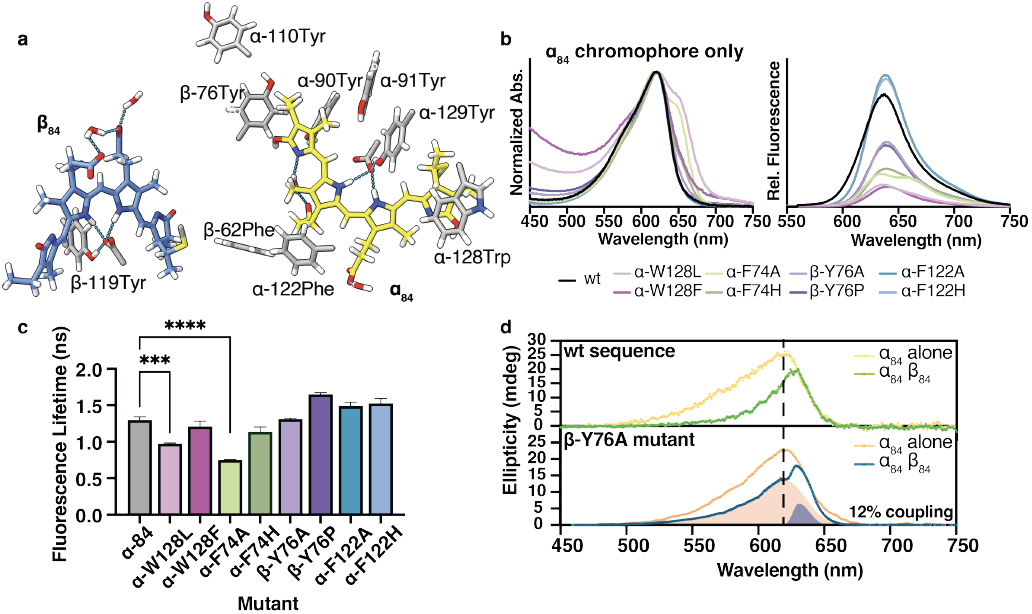
Examining the effects of the aromatic residues surrounding the α_84_/β_84_-coupled chromophore pair in CPC. **(a)** A highly conserved network of aromatic amino acids surrounds the central chromophores in CPC. **(b)** Plots are shown for the normalized absorption spectra and relative fluorescence intensities of the α_84_ chromophores and mutants. **(c)** Amplitude-averaged fluorescence lifetime measurements showed a significant reduction in the lifetimes of the α-W128L and α-F74A mutants (from an unpaired 2-tailed *t*-test). The β-Y76P, α-F122A, and α-F122H mutants were also found to have a statistically significant increase in fluorescence lifetime (p ≤ 0.01). **(d)** CD spectra of β-Y76A mutants known to possess full chromophorylation levels showed a reduced extent of coupling associated with the mutation. Split Gaussian decomposition estimated only 12% contribution by area of the sharp, red-shifted CD band associated with chromophore coupling.

Using the heterologous expression system, we mutated a few residues of interest in the aromatic network around the coupled α_84_/β_84_ pair, selecting residues α-74Phe, α-122Phe, α-128Trp, α-129Tyr, β-76Tyr, and β-119Tyr, which lie in close proximity to the chromophores and showed high sequence conservation across cyanobacterial species. In previous studies, mutation of specific aromatic residues surrounding the chromophore to an alanine led to reduced energy transfer to PSII, [34] but the systems biology approach leaves unanswered questions as to whether this mutation led to deleterious properties toward protein expression, chromophore attachment, or the excitonic interactions themselves.

We began by screening how the nearby mutations would affect the spectral properties of the α_84_ chromophore alone. Mutation of a tyrosine to a phenylalanine, α-Y129F and β-Y119F, therefore leading to removal of the hydrogen bonding motif of the tyrosine residues axial to the pigments, showed minimal effects on the energetics of the chromophores (Figure S9). In contrast, we found that introducing the Y-to-A mutation as previously studied *in vivo* showed no chromophore attachment. [34] This suggested that, rather than the elimination of the aromatic network being the sole cause of the disruption to energy flow from CPC complexes, the absence of the chromophore caused by disruption of the protein binding site is likely the main contributor to EET inhibition in the native organism.

Also of interest, we found mutations that led to much diminished fluorescence intensities even with full chromophorylation. The α-W128L mutant displayed this phenomenon and was accompanied by a red-shifted shoulder in the absorbance spectrum, even with only a single set of chromophores present (**Figure 7b**). This mutant also showed a significantly shorter fluorescence lifetime (**Figure 7c**), suggesting the role of this tryptophan as a sort of “tongue depressor” that rigidly holds the α_84_ chromophore into position. We hypothesize that elimination of this bulky aromatic residue leads to the accessibility of two major states of chromophore geometry, which we are currently investigating through *in silico* techniques.

In contrast, the α_84_-chromophorylated α-F122A and β-Y76A mutants showed similar energies, fluorescence intensities, and fluorescence lifetimes to the wild-type CPC containing just the α_84_ chromophore (**Figure 7b**,**c**). A difference in the response only appeared when we introduced the second set of chromophores at β_84_, which form the α_84_/β_84_ coupled pair as previously discussed. We expressed the α_84_β_84_ mutant CPCs and found reduced chromophorylation efficiencies in the α-F122A mutant (Figure S11); however, the β-Y76A mutant showed full chromophorylation at both the α_84_ and β_84_ positions, so we examined the chromophore excitonic interactions using CD spectroscopy.

We observed a diminished extent of chromophore coupling shown by the bimodal CD spectrum in **Figure 7d**, with two contributing bands corresponding to an excitonically coupled interaction with a maximum at 630 nm (dark blue) and a non-coupled band with a maximum around 620 nm (orange). We decomposed the CD spectrum of the β-Y76A mutant into asymmetric Gaussian bands, described in Table S3, and from the areas under the curves calculated the percent contribution of each. [35, 36] This gave rise to a 12% contribution of the 630 nm CD band, indicating that the β-Y76A mutation resulted in a lower extent of α_84_/β_84_ chromophore coupling. The normalized CD spectra given by each α_84_ CPC variant, the wild-type sequence or the β-Y76A mutant, were subtracted from the analogous α_84_β_84_ CPC CD signal to prepare difference spectra (Figure S12). The resulting S-shaped bands indicated weak electronic coupling between the α_84_ and β_84_ chromophores. [26, 27] We also analyzed the CD difference spectrum generated between the uncoupled α_84_ and β_155_ chromophores to benchmark these differences against non-coupled chromophores. Comparison of the amplitudes of the difference signals likewise indicated significantly diminished coupling strengths in the β-Y76A mutant, with a 37% decrease in the amplitude between wild-type and β-Y76A CPC (Figure S12).

Taken together, these results imply that this aromatic residue does not confer increased rigidity to enhance EET, as shown by the single-chromophore analysis of lifetimes and fluorescence intensities; instead, β-Y76 holds the chromophores in more optimal configurations that allow for critical chromophore interactions for energy transfer. The results from this mutational study and the interesting observed phenomena provide future opportunities for in-depth molecular dynamics and excited-state geometry calculations that could reveal how small changes affect the behavior of the chromophores *in silico*.

## CONCLUSION

We have recombinantly expressed a trimeric phycocyanin complex, and through the regiospecificity of each chromophore lyase, have selectively created minimal chromophore sets of CPCs to study the contributions of individual PCBs to the function of the overall complex. This approach provided a clear picture of the spectral line shapes and intrinsic lifetimes of each chromophore within its protein environment. The introduction of a quencher that can absorb the chromophore excitation energy at various positions within CPC confirmed the ability of the protein environment to tune the directionality of energy transport in this assembly. Further explorations of the role of key interactions with the pigments were explored, showing the deeper insight this approach can provide to understanding the impact of structure on the EET mechanisms in the phycobilisome.

In all, the power of this recombinant expression is the modularity with which we can examine different aspects of the chromophores in CPC. This allows us to understand in the simplest terms how a structural modification will directly affect the chromophore of interest before bringing in other interacting chromophores. This opens the door to studying a much wider breadth of mutations systematically to understand the sequence-structure-function relationship of the chromophores within the phycobiliproteins. Further exploration of the effect of the assembly state of CPC (monomeric, trimeric, or hexameric), and the effects that the linker proteins from the PBS have on the spectroscopic properties of CPC, is currently under investigation using this heterologous expression system.

## EXPERIMENTAL SECTION

### General Methods

Unless otherwise noted, all chemicals and solvents were of analytical grade and received from commercial sources. Water (dd-H_2_O) used in biological procedures and as a solvent was deionized to a resistivity of 18.2 MΩ⋅cm at 25 °C using a Barnstead NANOpure purification system (ThermoFisher, Waltham, MA). Duet^TM^ vectors were purchased from EMD Biosciences, Inc. (Burlington, MA) and gene blocks were purchased from Integrated DNA Technologies (Coralville, IA).

### Instrumental Analysis

#### Mass spectrometry

Electrospray ionization mass spectrometry (ESI-MS) of proteins and their bioconjugates was performed using an Agilent 1260 series liquid chromatography (LC) system that was connected in-line with an Agilent 6530 Q-TOF MS system (Santa Clara, CA). Protein samples were injected onto a Proswift RP-4H (monolithic phenyl, 1.0 mm × 50 mm, Dionex, Sunnyvale, CA) analytical column with a flow rate of 0.400 mL/min and a gradient of 5-100% MeCN + 0.1% (v/v) formic acid (FA) in dd-H_2_O + 0.1% FA over the course of 8 min. The total ion count chromatograms were extracted in the Agilent Mass Hunter Workstation software (Qualitative Analysis Version B 10.0, Build 10.0) and protein molecular weights were reconstructed using a maximum entropy deconvolution.

#### High Performance Liquid Chromatography (HPLC)

HPLC was performed on an Agilent 1100 Series HPLC System. Sample analysis for all HPLC experiments was achieved with an in-line diode array detector (DAD) and in-line fluorescence detector (FLD). Size exclusion chromatography (SEC) was performed using an AdvanceBio SEC 300 Å column (7.8 × 300 mm, 2.7 μm) (Agilent, Santa Clara, CA) at 1.0 mL/min using a mobile phase of 200 mM sodium acetate buffer, pH 5.5.

#### Steady-State Spectroscopy

UV-Vis absorbance measurements were conducted on a Cary UV-Vis 100 spectrophotometer (Agilent, Santa Clara, CA) and fluorescence emission measurements were conducted on a FluoroMax-4 fluorometer (HORIBA, Sunnyvale, CA). Protein concentration was determined by UV-Vis analysis on a Nanodrop 1000 instrument (ThermoFisher, Waltham, MA) by monitoring absorbance at 280 nm. The absorbance at visible wavelength (400-800 nm) of each line of *S. elongatus* PCC 7942 (WT, Δα-85 Δβ-154, and Δα-85 Δβ-83) was measured using a UV-3600 Plus Spectrophotometer with an integrating sphere (Shimadzu, Pleasanton, CA).

#### Circular Dichroism (CD) Spectroscopy

CD measurements were conducted using an Aviv 410 CD spectrophotometer (Biomedical Inc., Lakewood, NJ) with a 1 mm path quartz cuvette. All measurements were taken at 25 °C, recording from 400-700 at 1.0 nm intervals. Blank buffer samples in the absence of proteins were also recorded for baseline corrections.

#### Time-Resolved Fluorescence Spectroscopy

Fluorescence lifetime measurements were collected via time-correlated single photon counting (TCSPC) using a PicoQuant FluoTime FT-300 fluorometer (Berlin, Germany). The samples were transferred to a 1 mm path-length quartz cuvette and excited with a 520 nm PicoQuant pulsed diode laser, with an instrument response function of 150 ps as measured with a scattering LUDOX sample. The lifetime values result from mono- or bi-exponential deconvolution fitting using PicoQuant FluoFit software version 4.6.6.0, with χ^2^ < 1.1 for all measurements.

### Molecular Biology Techniques

#### Construction of Expression Plasmids

The highest copy number plasmid, pRSFDuet-1 containing the pRSF1030 replicon, was used for genes for the α and β chain of CPC. The vector was digested with NcoI and AflII, then the digested fragment was purified on a 1% agarose gel. Gibson assembly was used to ligate a gene block containing CpcA (Uniprot ID: P13530), another T7 promoter, LacO, and rbs before the CpcS gene (Uniprot ID: Q8YZ70). This plasmid was then digested with NdeI and AvrII, purified, then ligated with the gene block encoding for CpcB (Uniprot ID: P06539). The genes encoding for the phycocyanobilin production enzymes were encoded in a lower copy number plasmid, pETDuet-1 with the ColE1 replicon in a similar fashion, taking the genes for BHo1 (Uniprot ID: Q8DLW1) and PcyA (Uniprot ID: Q8KPS9). In the third vector, the genes for CpcE (Uniprot ID: Q44115), CpcF (Uniprot ID: Q44116), and CpcT (Uniprot ID: Q31Q65) were inserted into pCDFDuet-1 containing the CloDF13 origin of replication.

#### Protein Expression and Purification

BL21 Star (DE3) competent cells were transformed with the three plasmids containing the CPC complexes described above. Colonies were selected for inoculation in Terrific Broth with 100 μg/L carbenicillin and 50 μg/L of both kanamycin and streptomycin at 37 °C. When cultures reached an optical density of 0.6-0.8 at 600 nm, IPTG was added to a final concentration of 1 mM. After growing for 20 h at 20 °C, the cells were harvested by centrifugation (8000 rpm, 30 min). Cells were resuspended in 20 mL of lysis buffer (20 mM triethanolamine (TEOA), pH 7.2) supplemented with 2500 units of Benzonase® nuclease and 2 mM MgCl_2_. Cells were lysed by sonication with a 2 s on, 4 s off cycle for a total of 10 min on using a standard disruptor horn at 70% amplitude (Branson Ultrasonics, Danbury, CT). The resulting lysate was cleared at 14000 rpm for 30 min. A saturated solution of ammonium sulfate was added to the supernatant to reach the final concentration of 30%. The mixture was rotated for 10 min at 4 °C to allow the complete protein precipitation. The precipitated protein was then collected at 11000 rpm for 30 min and then resuspended in lysis buffer. The resulting protein solution was dialyzed against the lysis buffer to remove the residual ammonium sulfate before loading onto a DEAE column and purified with a 0-600 mM NaCl gradient elution in lysis buffer. Purity was confirmed by SDS-PAGE and ESI-TOF-MS.

#### Gel Analyses

Sodium dodecyl sulfate-polyacrylamide gel electrophoresis (SDS-PAGE) was carried out in a Mini cell tank apparatus (Life Technologies, Carlsbad, CA) using Nu-PAGE Novex 4-12% Bis-Tris Protein Gels (Life Technologies). The sample and electrode buffers were prepared according to the suggestions of the manufacturer. All protein electrophoresis samples were heated for 10 min at 95 °C in the presence of 1,4-dithiothreitol (DTT), 10% (w/v) SDS, 0.05% (w/v) bromophenol blue. Gels were run for 30 min at 200 V to separate the bands. Commercially available markers were applied to at least one lane of each gel for the assignment of apparent molecular masses. Native PAGE was performed using NativePAGE 4-16% Bis Tris Protein Gels (Life Technologies) and 200 mM NaOAc, pH 5.5. Samples were mixed with 80% glycerol at a sample/glycerol ratio of 1:1 and allowed to settle in wells for 10 min prior to applying voltage. Native gels were placed on ice and run for 2 h at 25 V to separate the bands. Visualization of protein bands was accomplished by staining with Coomassie Brilliant Blue R-250 (Bio-Rad, Hercules, CA) and imaged on a Gel Doc EZ Imager (Bio-Rad).

#### Labeling CPC with Atto MB2 Maleimide

Atto MB2 maleimide was purchased from ATTO-TEC (Seigen, Germany) and dissolved in DMSO to make a 1 mM stock solution. To 500 μM of CPC in 20 mM TEOA, pH 7.5 was added 2 equiv. of the maleimide functionalized quencher. The reaction mixture was briefly agitated and then incubated at 4 °C for at least 4 h. The crude reactions were then purified with a NAP-5 Sephadex G-25 column (GE Healthcare, USA) to remove the excess small molecules.

#### Site-directed mutagenesis of CPC in cyanobacteria

The two DNA fragments for the cpcB1-cpcA1 cluster were synthesized with C-to-V mutations: i) cpcB1(C83V)-cpcA1(C85V) and ii) cpcB1(C154V)-cpcA1(C85V); and cloned each of them into the pAM2991 plasmid (a gift from Susan Golden through Addgene), which is designed to insert the mutated cpcB1-cpcA1 cluster into the neutral site I (NS1) with a *trc* promoter (P_trc_) on the 5’ flanking region and a spectinomycin-resistant gene cassette on the 3’ flanking region via homologous recombination. [37] After generating each of the overexpressed lines, we knocked out endogenous genes for the cpcB1-cpcA1 and the cpcB2-cpcA2 clusters by inserting a gentamicin-resistant gene cassette and a kanamycin-resistant gene cassette, respectively, through homologous recombination using the 1-kbp flanking regions.

The transformation of *S. elongatus* PCC 7942 was done according to the previous literature. [38] In brief, the WT was grown in BG11 liquid media under continuous light (fluorescence bulbs) at ∼100 µmol photons m^-2^ s^-1^. A 50 mL culture with a cell density of 0.5 (OD_730_) was centrifuged at 1,300 × *g* for 5 min, and the cell pellet was washed once with 10 mL of 10 mM NaCl solution. After washing, the cell pellet was resuspended in 1 mL of BG11 liquid media. Next, 200 µL aliquots of the cell suspension were transferred to 1.5 mL tubes and mixed with 1 µg of the DNA plasmid described above. Transformation occurred by incubating the cells in dark conditions overnight with gentle agitation at room temperature. Different antibiotics, including spectinomycin, kanamycin, and gentamicin, were used for selection on BG11 plates, with concentrations of 20 µg/mL, 10 µg/mL, and 5 µg/mL, respectively.

#### Growth measurements

The WT and two mutant lines (Δα_85_Δβ_154_ and Δα_85_Δβ_83_) of *S. elongatus* PCC 7942 were grown in BG11 liquid media (50 mL) with spectinomycin (20 µg/mL), kanamycin (10 µg/mL), and gentamicin (5 µg/mL) in a sterile 250-mL polypropylene beaker covered with a polystyrene Petri dish bottom for evenly distributed light illumination and shaken at 120 rpm at 23 °C under continuous light (fluorescent bulbs) at 3, 40, or 285 µmol photons m^-2^ s^-1^. The OD_730_ was measured using a SpectraMax ABS Microplate Reader (Molecular Devices, San Jose, CA) for the growth curve with three biological replicates.

#### PBS isolation

The isolation of PBS from *S. elongatus* PCC 7942 was done according to the previously described method. [39] The WT and two mutant lines (Δα_85_Δβ_154_ and Δα_85_Δβ_83_) were grown in BG11 liquid media (100 mL) with spectinomycin (20 µg/mL), kanamycin (10 µg/mL), and gentamicin (5 µg/mL) in a 500-mL flask shaken at 120 rpm at 23 °C under continuous light (fluorescent bulbs) at 40 µmol photons m^-2^ s^-1^. The culture was centrifuged at 3,000 × *g* for 10 min at 23

°C. The cell pellet was washed with 10 mL of 0.6 M potassium phosphate (KP) buffer (pH 7.0) and centrifuged at 3,000 × *g* for 10 min at 23 °C. The cell pellet was stored at -20 °C until used for ultracentrifugation. The thawed cell pellet was washed twice with KP buffer in the same way as above. The washed cells were resuspended with 0.75 M KP buffer (the cell density was adjusted to the OD_730_ of 0.5-0.8 for the 1:100 diluted sample). The resuspended cells were lysed with acid-washed 425-600 µm glass beads (the volume of glass beads was half of the resuspended cells) by vortexing for 1 min three times with a 1 min interval between each. After removing glass beads, the cell lysate was solubilized with the final concentration of 2% (w/v) Triton X-100 for 30 min at room temperature with gentle mixing every 5 min. Then, the sample was centrifuged at 20,800 × *g* for 20 min at 23 °C. The supernatant was loaded onto sucrose gradients, containing 10 to 50% (w/v) sucrose with 0.75 M KP buffer, and the phycobilisome bands were separated at 154,300 × *g* for 18 h at 18 °C (SW 41 Ti, Beckman Coulter).

### Computational Protocols

#### Computational Modeling of Phycocyanin

To investigate the protein environment around each chromophore, we processed the hexameric CPC structure from *Synechococcus elongatus* sp. PCC 7942 from the published crystal structure (PDB ID: 4H0M) to include hydrogen atoms with protonation states at pH 7.4 using the protein preparation module in Maestro (version 2024-4), while ensuring the protonation of the PCB chromophore and deprotonation of the aspartate residue axial to the chromophore plane and optimization of the hydrogen bonds. The N-methylated asparagine (MEN) was mutated to an asparagine to better reflect the CPC used in *in vitro* experiments.

#### Molecular Dynamics (MD) Simulations

The Schrödinger Maestro package (version 2024-4) was used for structural preparation and molecular dynamics simulations. The Desmond system builder was used to solvate the double disk structure with water molecules described using the TIP3P model in a rhombic dodecahedron *xy*-square box with periodic boundaries of 15 Å from the protein and neutralized with sodium ions with the addition of 150 mM NaCl in an OPLS5 force field. Simulations of each complex were completed in triplicate, using a RESPA-based Multigrator integration scheme and a timestep of 2 fs unless otherwise specified. Force constants for restraints were set to 50 kcal⋅Å^-1^, and NPT simulations were conducted at 1 atm. The van der Waals and short-range electrostatic interactions cutoff was set to 9 Å, and long-range electrostatics were calculated every 6 fs for the NPT simulations and every 3 fs for the NVT equilibration stages.

The first four stages of equilibration used a Langevin thermostat with a 0.1 ps relaxation time and a Langevin barostat with a 50 ps relaxation time when applicable. The first stage was an NVT equilibration with Brownian dynamics, a 1 fs timestep, restraints on heavy solute atoms, and was run at 10 K for 100 ps. The second stage was an NVT equilibration run with Langevin dynamics with a 1 fs timestep and the same restraints at 10 K for 50 ps. The third and fourth stages were NPT equilibrations with heavy solute atoms restrained and run for 500 ps at 10 K, then 300 K. The fifth and last stage was an NPT equilibration with no restraints at 300 K for 500 ps and used a Langevin thermostat and barostat with a 0.1 ps and 2.0 ps relaxation time, respectively. The production simulation was run in the NPT ensemble using the Martyna-Tobias-Klein barostat and Nose-Hoover thermostat with a 1 ps coupling time at 300 K for 150 ns.

Convergence across all three replicates was determined by measuring both the root mean square deviation (RMSD) of the protein backbone as well as the cosine content of the first principal component (PC) of the simulation. Cosine content measurement was performed in addition to RMSD measurements as there were some simulations for which the RMSD plot did not have a clear plateau and was thus insufficient for determining equilibration. MDAnalysis was used for principal component analysis (PCA), a mathematical technique that in this case reduces all the motions in an MD simulation to key movements in the trajectory. [20] The cosine content of the first principal component can then be used as a gauge for system equilibration, as the closer the value is to 1, the closer it resembles a perfect cosine curve and random diffusion. Cosine content measurement was conducted as described by Hess, and values under 0.7 are used to indicate sufficiently sampled, equilibrated systems. [21]

#### TD-DFT Calculations

Vertical excitation energies, oscillator strengths, and natural transition orbitals (NTOs) with ground state (S_0_) and excited state (S_1_) minimum energy geometries were calculated in Jaguar using TD-DFT at the B3LYP-D3/6-31G+** level, with aqueous solvation effects from a standard polarizable continuum model. The chromophore geometries were taken directly from the crystal structure with added hydrogen atoms to match the expected protonation state of the chromophore. The side chains of hydrogen bonded residues were also included, as well as water molecules present from the crystal structure.

#### Förster Rate Calculations

Due to the expected interchromophore distances in this system, any energy transfer between chromophores in CPC would be due to FRET. Thus, we set out to calculate the rate *k*_*ij*_ between each site *i* and *j* within a CPC trimer assembly using Förster theory. [40]

Interchromophore distances were measured based on their center of mass in ChimeraX, version 1.9. The output transition dipole moments (TDMs) of the S_0_ → S_1_ transitions with the highest oscillator strength, reflecting the transition with the highest probability of occurring, were used then in then calculating the dipole orientation factor between each chromophore

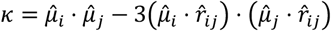

where 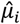 and 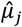 are the unit TDM vectors of the chromophore *i* and *j* and 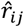 is the unit vector point from site *i* to site *j*.

The Scholes group has shown that a dipole-dipole approximation in adequate for describing the strength of electronic coupling when the separation between chromophores is greater than 20 Å. [41, 42] Therefore, we approximate V_ij_ as

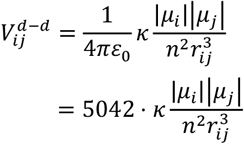

where a dielectric constant of *n*^*2*^ = 2 is used for protein environments. [41, 42]

Last, J is the overlap integral of the acceptor’s molar absorptivity spectrum 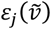 and the donor’s area-normalized fluorescence spectrum 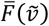:

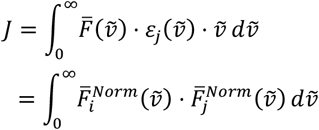

which, when evaluated over energy in wavenumbers (cm^-1^) has units of cm^6^⋅mol^-1^.

The spectral overlap is dependent on the line shapes of the acceptor and donor spectra, so based on the 25 calculated excited states of the PCB chromophore calculated in Jaguar (in cm^-1^), the absorption spectra were fitted with Gaussian broadening with half bandwidth of 400 cm^-1^. [43, 44] The donor emission line shapes were made by mirroring the corresponding absorption spectrum and red-shifting by the experimentally reported Stokes shift.

Finally, the energy rate can then be approximated as

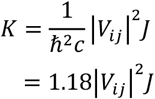

where K is given in units of ns^-1^. [45]

#### Transfer Efficiency Calculations

To calculate the quenching efficiency in the CPC-quencher systems, the following expression was used

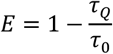

where E indicates energy transfer efficiency from PCB to the MB quencher.

## ABBREVIATIONS

CPC: C-phycocyanin
PCB: phycocyanobilin
PBS: phycobilisome
EET: excitation energy transfer
CD: circular dichroism
TDM: transition dipole moment
FRET: Förster resonant energy transfer.

## ASSOCIATED CONTENT

### Supporting Information

The Supporting Information is available free of charge on the ACS Publications website.

Mass spectra of purified proteins and steady-state UV-Vis spectra (PDF)

## AUTHOR INFORMATION

### Author Contributions

The manuscript was written through contributions of all authors. / All authors have given approval to the final version of the manuscript.

## ACKNOWLEDGMENT

We thank Dr. Kathleen Durkin, Dr. Azhagiya Singam, and the Molecular Graphics and Computing Facility (MGCF) in the UC Berkeley College of Chemistry for the computational resources supported in part by NIH S10OD023532.

---

For Table of Contents only.

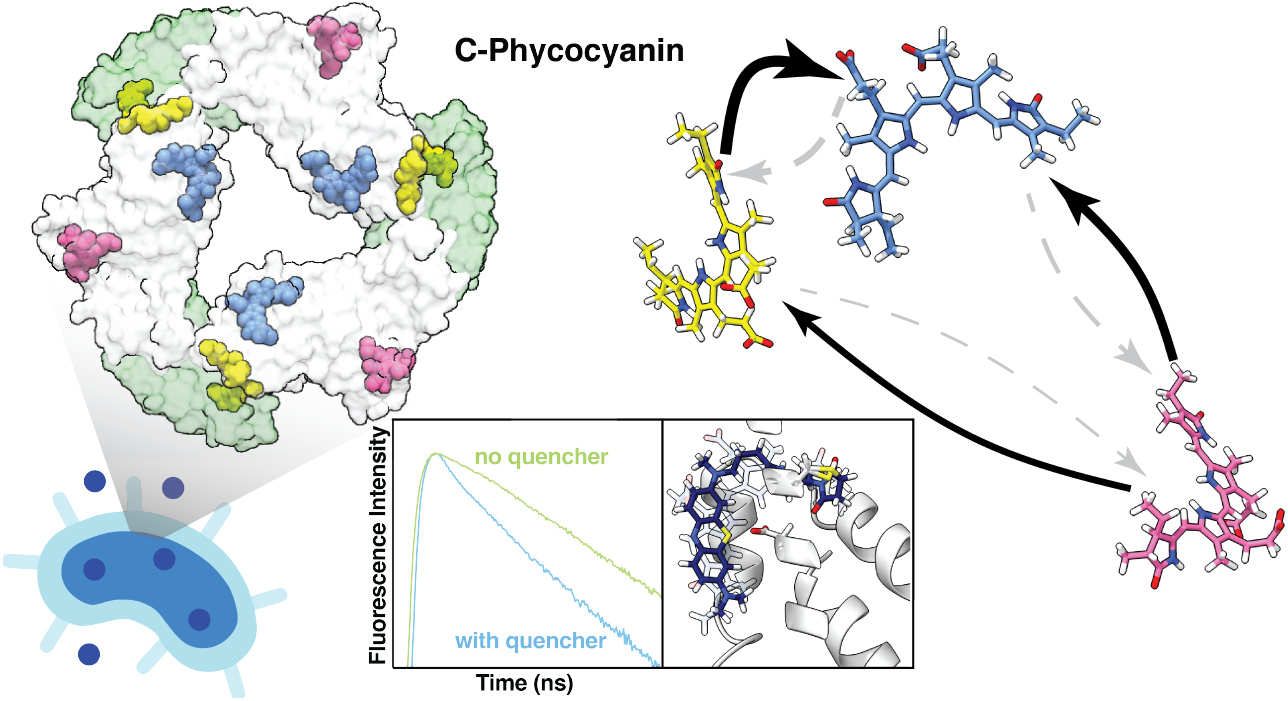

