## Supplementary Information for "Biosynthesis of minimal C-phycocyanin chromophore assemblies in *E. coli* provides a platform to dissect protein-mediated tuning of exciton transfer"

**Table of Contents**

**Scheme S1.** Biosynthetic pathway of PCB from heme. 2
**Figure S1.** Expression of CPC using no lyase enzymes. 3
**Figure S2.** Expression of mutants with no chromophore attachment. 3
**Figure S3.** Root-mean-square deviation (RMSD) of CPC backbones from MD simulations. 4
**Table S1.** Cosine content of the first principal component of CPC trimer complexes. 5
**Figure S4.** Structures of CPC variants colored by the per-residue RMSF. 6
**Figure S5.** Spectral predictions of the PCB chromophores as calculated by TD-DFT. 7
**Figure S6.** Spectral overlap of the Atto MB2 quencher with the CPC spectrum. 7
**Figure S7.** ESI-MS of ATTO-MB2-maleimide labeled CPC complexes. 8
**Table S2.** Förster approximations of electronic coupling strengths and energy transfer rates. 9
**Figure S8.** Transition dipole moments of quencher used to estimate electronic coupling. 9
**Figure S9.** ESI-MS of CPC with mutations to the axial tyrosine residues. 10
**Figure S10.** ESI-MS of CPC containing mutations with $\alpha_{84}$ chromophorylation. 11
**Figure S11.** ESI-MS of CPC containing mutations with $\alpha_{84}$ and $\beta_{84}$ chromophorylation. 12
**Table S3.** Gaussian decomposition of component peaks in β-Y76A mutant CD spectrum. 12
**Figure S12.** CD difference spectra between double-chromophore CPC sets and $a_{84}$ CPC. 13

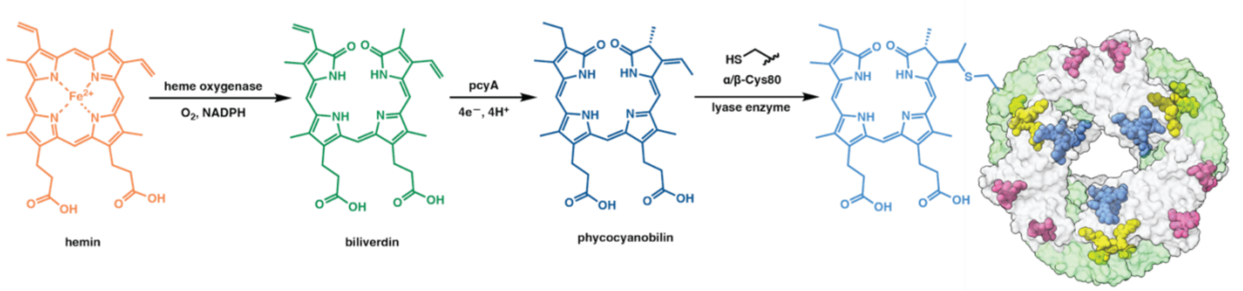


**Scheme S1.** Biosynthesis of chromophorylated CPC in *E. coli* converts heme into PCB by oxidative cleavage of the hemin methyne with heme oxygenase 1 (HO1) to form the linear tetrapyrrole biliverdin Ixα. The biliverdin then undergoes a 4-electron reduction to form the chromophore, phycocyanobilin (PCB). Three sets of lyase enzymes are then responsible for regiospecifically attaching PCB to cysteine residues at α-84, β-84, or β-155 in CPC by a Michael-like addition to form a thioether linkage.

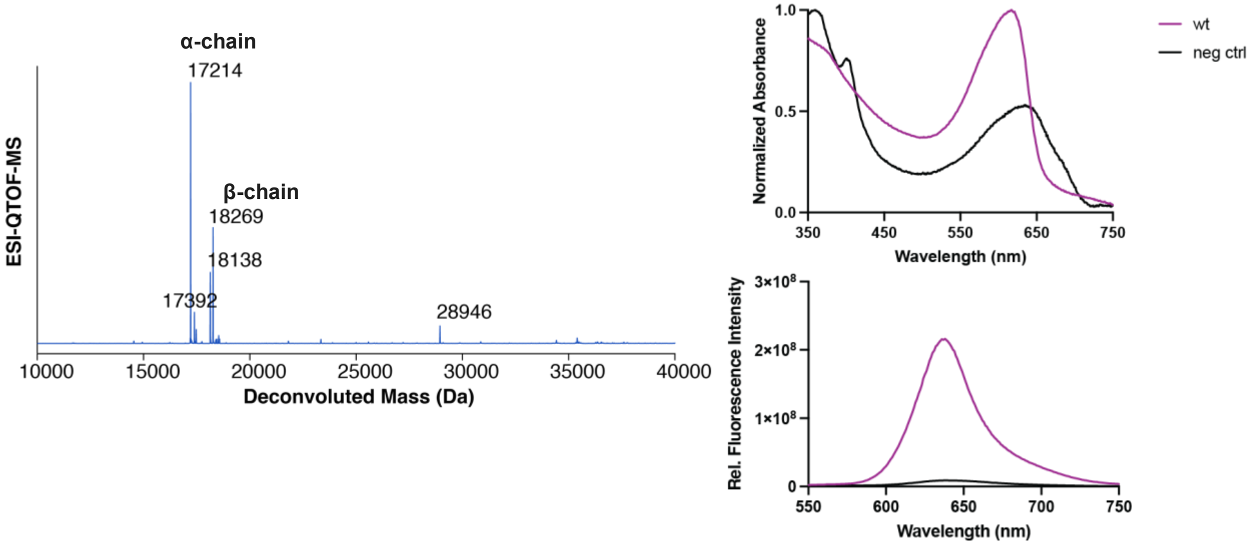


**Figure S1.** Expression of CPC in bacteria containing no lyase enzymes shows that there is minimal effect of nonspecific chromophore binding on the overall spectral properties of the expressed protein system. In addition, this experiment confirms that the free PCB chromophores in *E. coli* are not brightly fluorescent. Spectra were obtained following ammonium sulfate precipitation following lysis of *E. coli*.

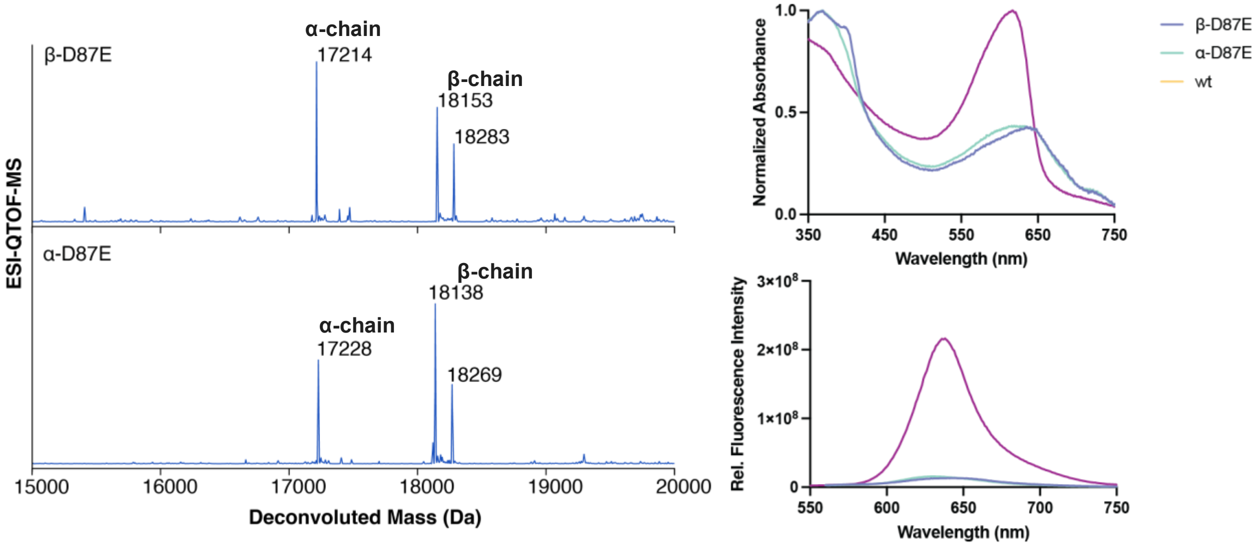


**Figure S2.** Mutations in the chromophore binding site prevent the covalent addition of the chromophore, showing that the lyases themselves do not contribute to the overall fluorescence signal. Spectra were obtained following ammonium sulfate precipitation following lysis of *E. coli*.

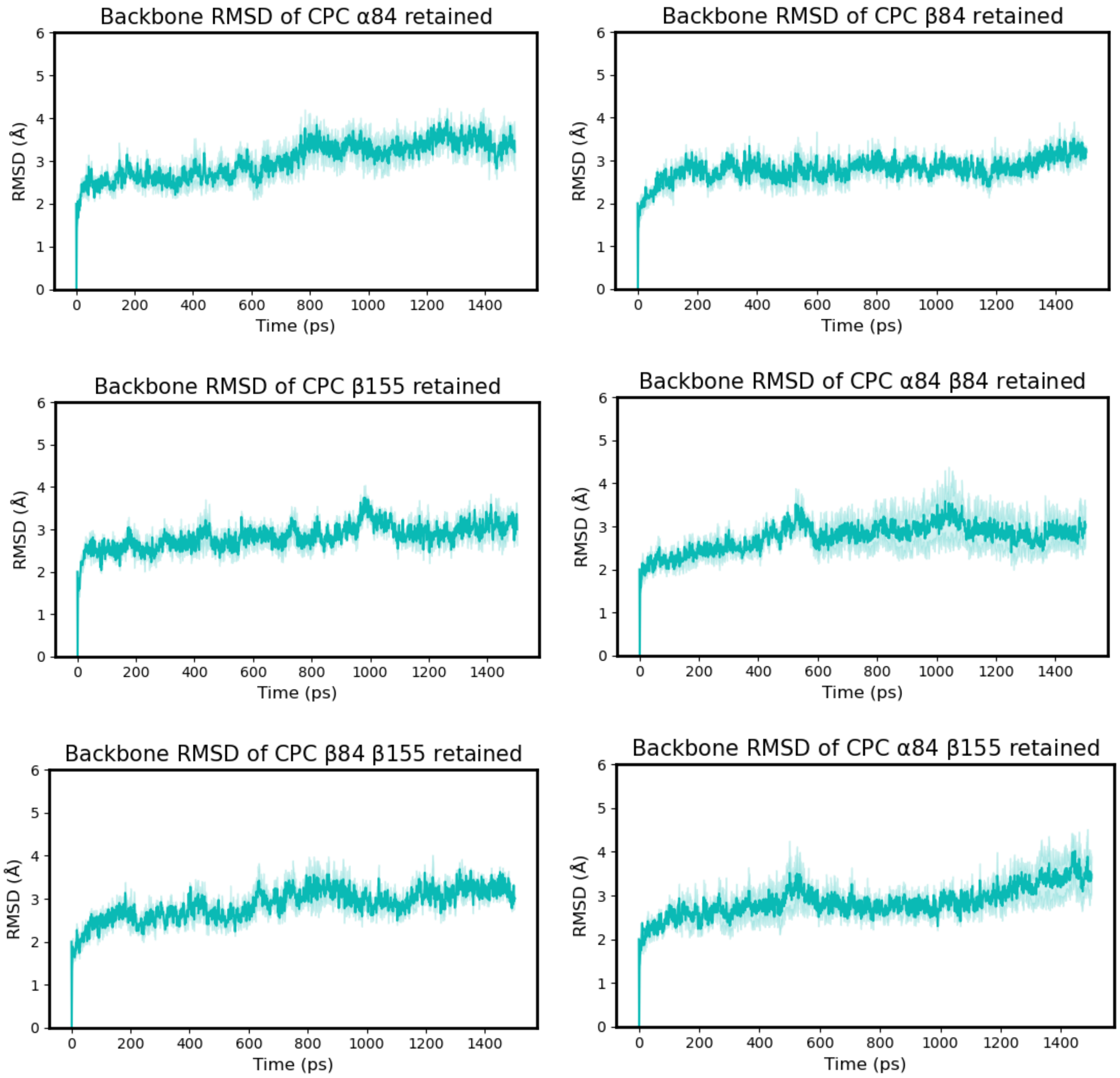


**Figure S3.** Root-mean-square deviation (RMSD) of Cα over the course of a 150 ns MD simulation of the trimeric CPC protein assembly systematically lacking sets of chromophores shows equilibration of the protein backbone during the simulation.

**Table S1.** Average radii of gyration and cosine content of the first principal component of CPC trimer complexes.

| System | Avg. radius of gyration (Å) | St. dev. (radius of gyration) | Avg. cosine content of PC 1[a] | Std. dev. (cosine content) |
|---|---|---|---|---|
| WT | 38.97 | 0.25 | 0.481 | 0.309 |
| α-C84V | 39.00 | 0.30 | 0.573 | 0.141 |
| β-C84V | 38.98 | 0.26 | 0.650 | 0.072 |
| β-C155V | 38.84 | 0.24 | 0.632 | 0.104 |
| α-C84V β-C84V | 39.17 | 0.25 | 0.736 | 0.053 |
| aC84V β-C155V | 39.05 | 0.19 | 0.513 | 0.270 |
| β-C84V β-C155V | 38.98 | 0.25 | 0.617 | 0.043 |
| α-C84V β-C84V β-C155V | 39.15 | 0.21 | 0.449 | 0.220 |

[a] Radius of gyration and cosine content computed using MDAnalysis. For cosine content, a value of 1 represents a perfect cosine wave and a value of 0 represents no cosine character.

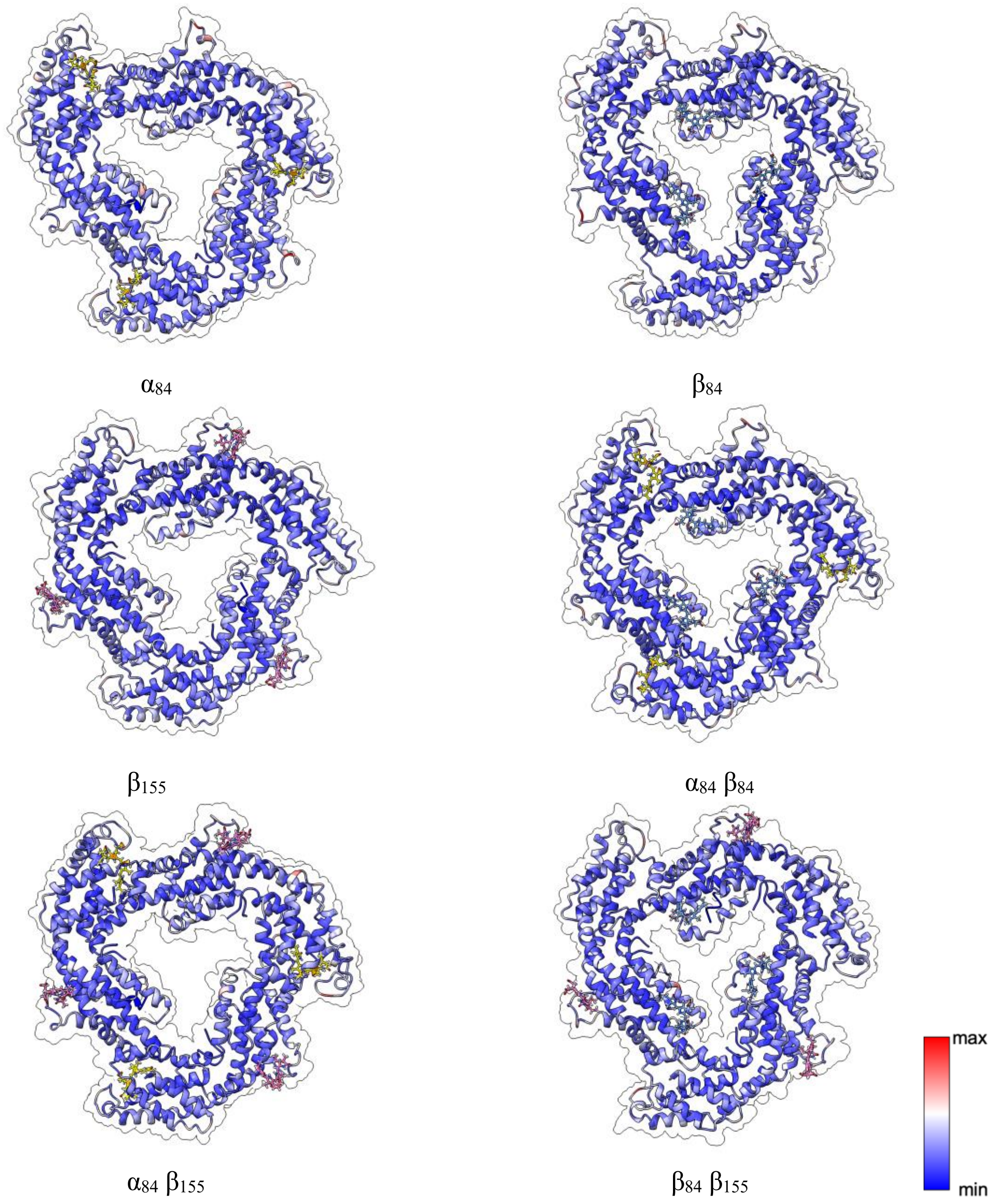


**Figure S4.** Structures of CPC variants colored by the per-residue RMSF of the α-carbons.

a

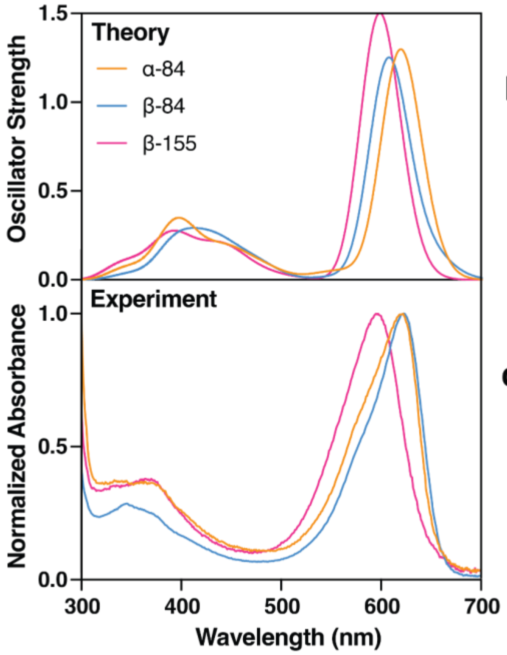


b

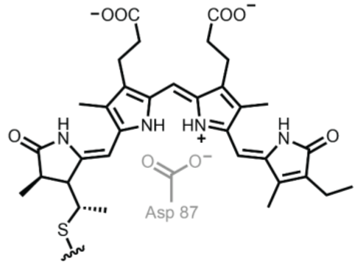


c

| CPC chromophore | Expt. (nm) | PCB Calc. (nm) | PCB-H⁺ Calc. (nm) |
| --- | --- | --- | --- |
| **α-84** | 622 | 591 | 619 |
| **β-84** | 623 | 585 | 608 |
| **β-155** | 597 | 562 | 598 |

**Figure S5.** Spectral predictions of the PCB chromophores taken from each position within CPC. Geometries from the crystal structure (PDB ID: 4H0M) and calculated using TD-DFT. Chromophore protonation was required to match the experimentally obtained spectra.

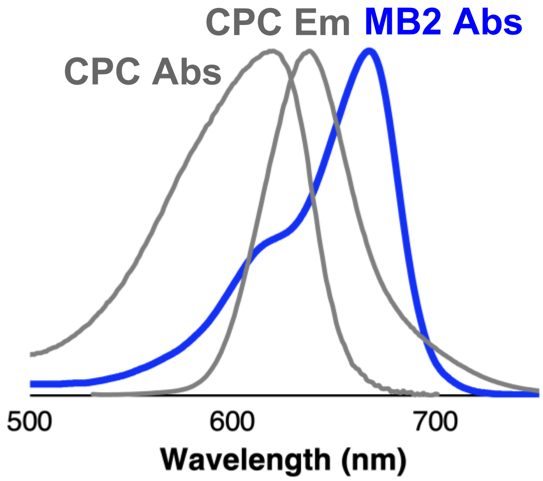


**Figure S6.** Spectral line shapes of the PCB chromophores within CPC (grey) overlaid with the absorbance trace of the quencher (Atto MB2, $\lambda_{max,\ Abs}$ = 668 nm, $\Phi_F$ < 0.04). This comparison showed appreciable spectral overlap between the acceptor and donor chromophores.

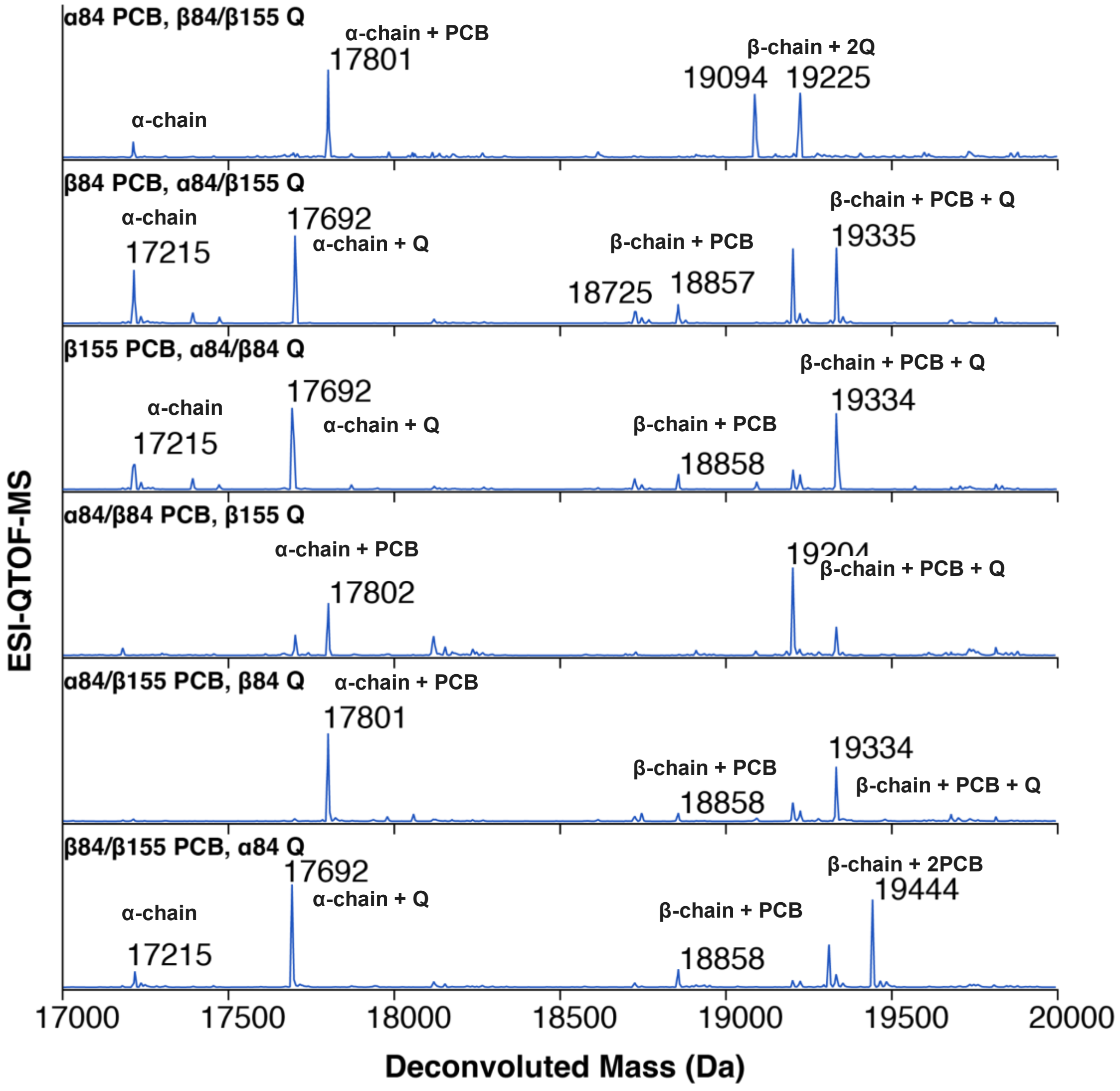


**Figure S7.** ESI-MS of ATTO-MB2-maleimide labeled CPC complexes.

| D-A pair | R (Å) | κ | \|V\| ($cm^{-1}$) | J (cm) | K ($ns^{-1}$) | 1/K (ps) | $1/K_{expt}$ (ps) |
|---|---|---|---|---|---|---|---|
| $\beta^1$-155 → $\alpha^1$-84 | 38.81 | 0.04 | 0.709 | 6.12E-4 | 0.31 | 3251 | >500 |
| $\alpha^3$-84 → $\beta^1$-84 | 20.31 | 1.41 | 113.34 | 3.46E-4 | 1094 | 0.91 | 0.1 – 1.0 |
| $\alpha^1$-84 → $\beta^1$-84 | 50.91 | 1.74 | 4.19 | 4.45E-4 | 7.80 | 128.2 | 200 ± 70 |
| $\beta^1$-84 → $\beta^2$-84 | 36.10 | -0.85 | 7.69 | 5.21E-4 | 30.80 | 32.47 | 45 |
| $\beta^1$-155 → $\beta^1$-84 | 35.09 | -0.57 | 11.79 | 5.58E-4 | 15.97 | 62.62 | 45 ± 15 |

**Table S2.** Estimated electronic coupling strengths and energy transfer rates from Förster approximations.

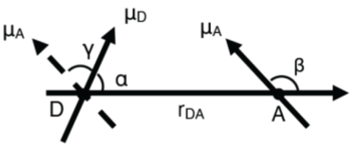


| Interactions | R (Å) | γ (°) | α (°) | β (°) | κ | V ($cm^{-1}$) |
|---|---|---|---|---|---|---|
| a-84 PCB to b-84 PCB | 20.83 | 116.8 | 32.0 | 138.4 | 1.451 | 141.09 |
| a-84 PCB to b-84 Q | 20.78 | 89.7 | 33.7 | 123.1 | 1.366 | 118.63 |
| b-84 PCB to a-84 Q | 16.51 | 86.9 | 107.3 | 140.3 | 0.632 | 101.38 |

**Figure S8.** Transition dipole moment analysis of quencher molecules. Data acquired from ligand docking of the quencher were used to estimate electronic coupling strengths. Förster theory analysis showed coupling strengths of similar magnitude of quenchers positioned at either α-84 or β-84.

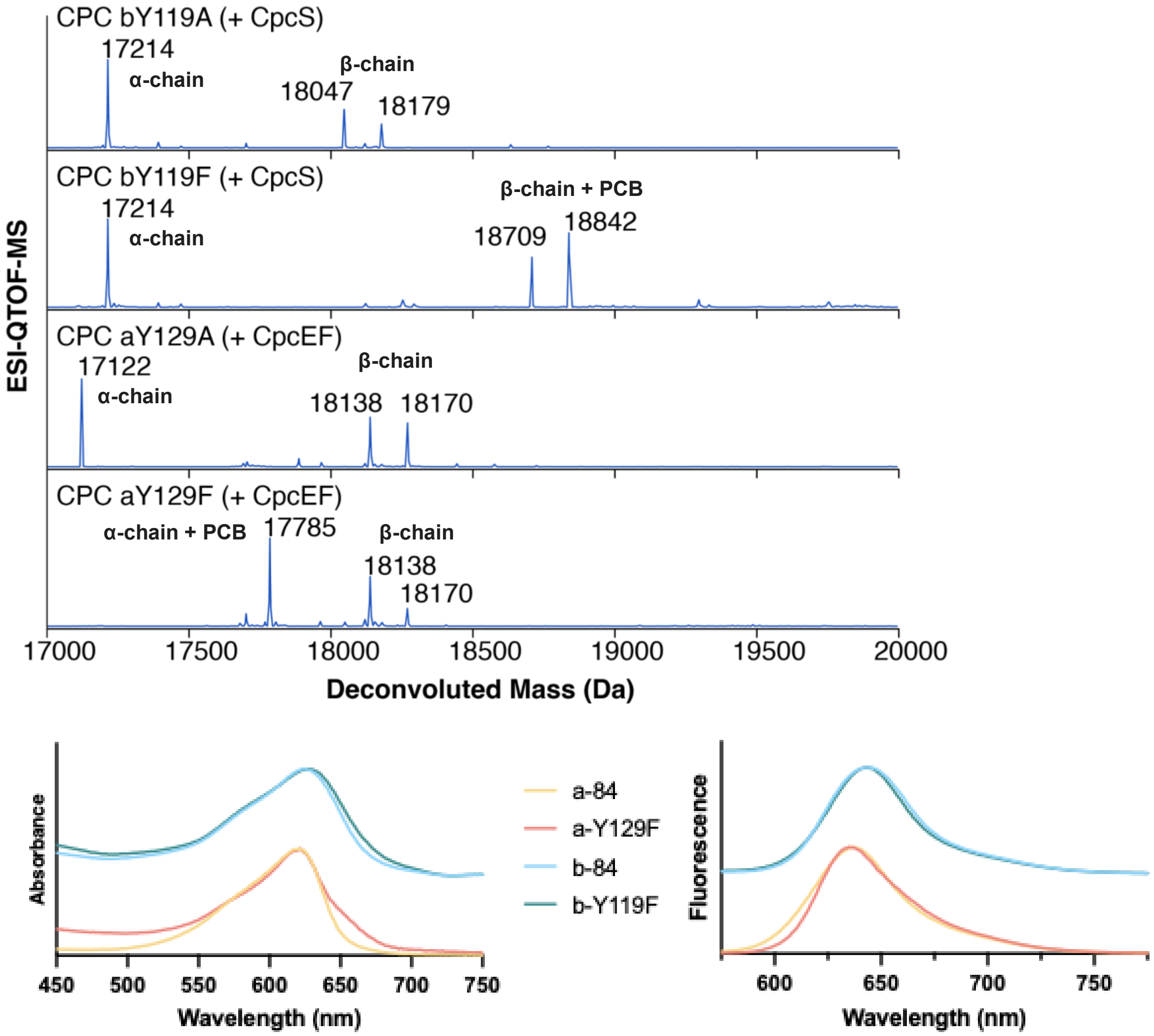


**Figure S9.** ESI-MS of CPC with mutations replacing the axial tyrosine residues near the $\alpha_{84}$ (Y129) and $\beta_{84}$ (Y119) chromophores.

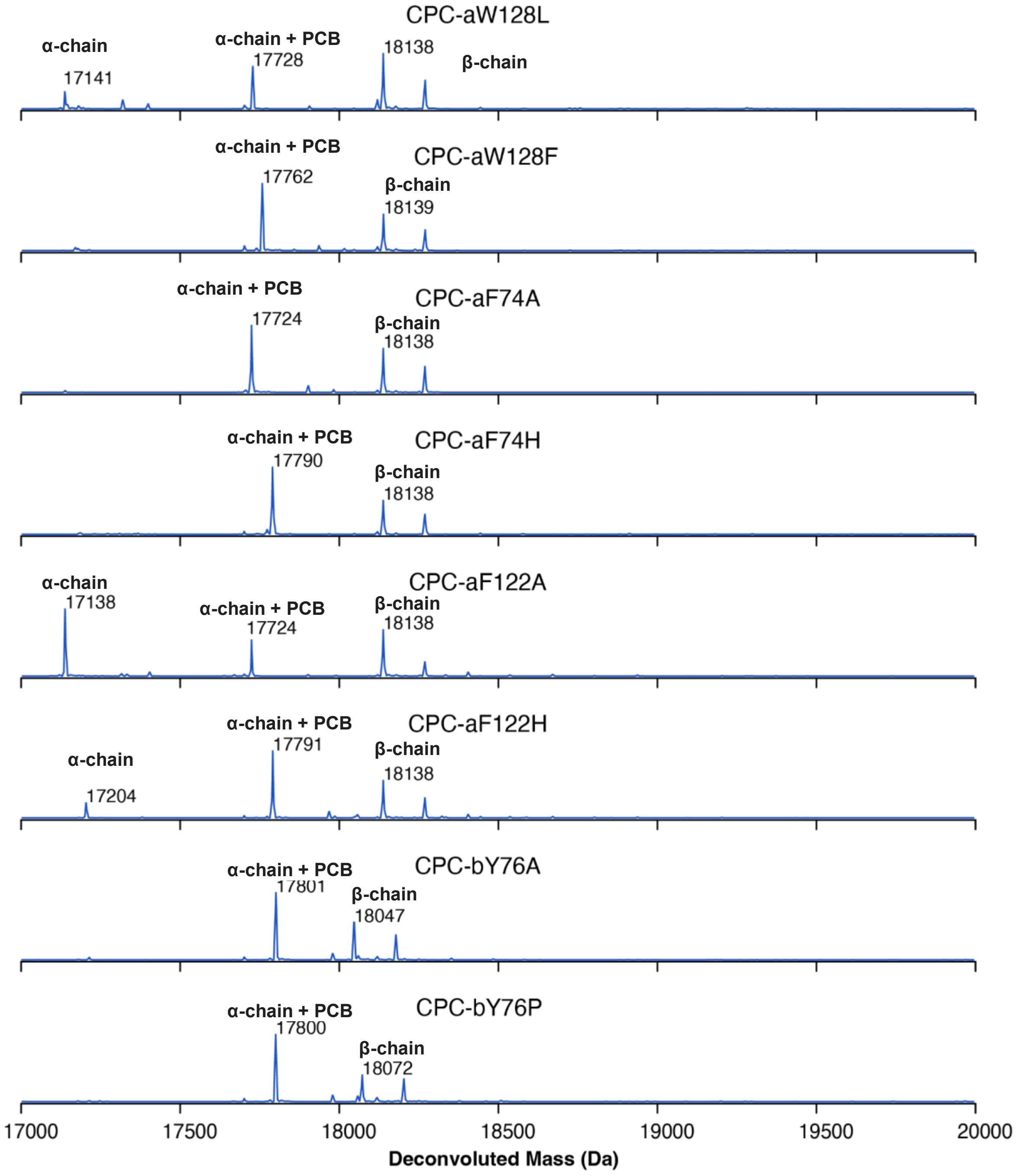


**Figure S10.** ESI-MS of CPC containing mutations replacing aromatic residues near the $\alpha_{84}$ chromophore. Only the CpcE/F lyase is incorporated to selectively chromophorylate only at $\alpha_{84}$.

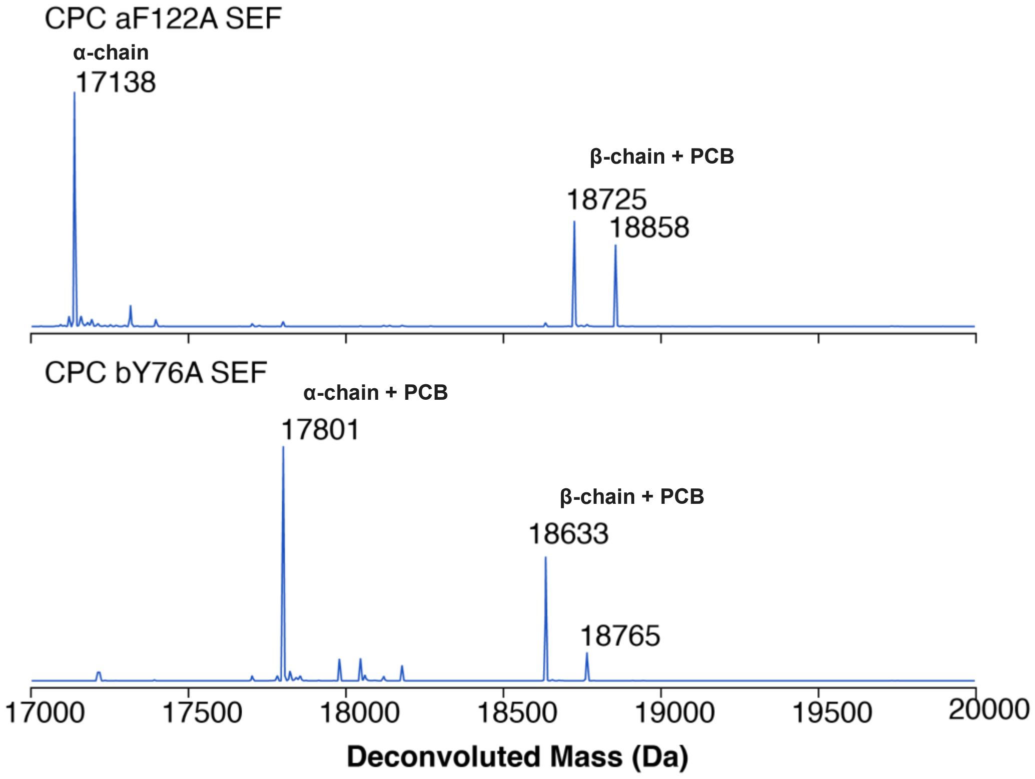


**Figure S11.** ESI-MS of CPC containing mutations to aromatic residues near the $\alpha_{84}$ chromophore with both $\alpha_{84}$ and $\beta_{84}$ chromophorylated.

**Table S3.** Gaussian band parameters for the decomposition of CD spectrum of β-Y76A $\alpha_{84}\beta_{84}$ mutant.

| Parameter | Peak 1 | Peak 2 |
|---|---|---|
| $\lambda_{max}$ | 621.5 | 631.5 |
| FWHM | 47.5 | 14.9 |
| HWHM | 29.3, 18.2 | 5.1, 9.8 |
| $\theta_{max}$ | 13.73 | 6.69 |
| area | 789.9 | 107.8 |
| %area | 88.0 | 12.0 |

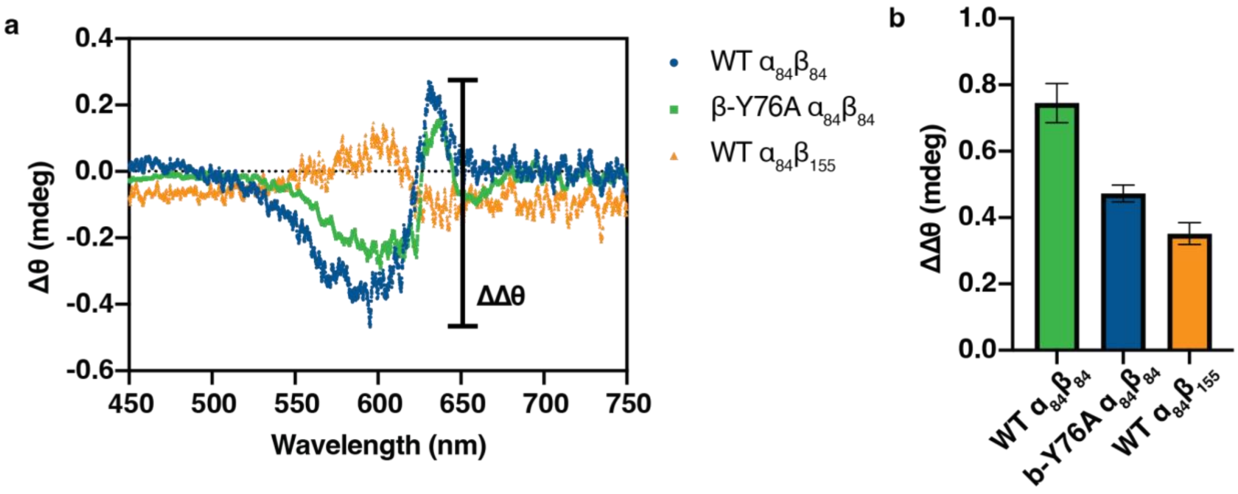


**Figure S12. (a)** CD difference spectra subtracting the a84-alone CD signal from CD signal of double-chromophore CPC sets reveals extent of electronic coupling between chromophores. **(b)** The splitting amplitude ΔΔθ decreases with decreasing coupling strengths.
